# Can two wrongs make a right? F508del-CFTR ion channel rescue by second-site mutations in its transmembrane domains

**DOI:** 10.1101/2021.09.07.459271

**Authors:** Stella Prins, Valentina Corradi, David N. Sheppard, D. Peter Tieleman, Paola Vergani

## Abstract

Deletion of phenylalanine 508 (F508del), in the cystic fibrosis transmembrane conductance regulator (CFTR) anion channel, is the most common cause of cystic fibrosis (CF). F508 is located on nucleotide-binding domain 1 (NBD1) in contact with cytosolic extensions of transmembrane helices, in particular intracellular loop 4 (ICL4). We carried out a mutagenesis scan of ICL4 by introducing five or six second-site mutations at eleven positions in *cis* with F508del, and quantifying changes in membrane proximity and ion-channel function of CFTR. The scan strongly validated the effectiveness of R1070W at rescuing F508del defects. Molecular dynamics simulations highlighted two features characterizing the ICL4/NBD1 interface of F508del/R1070W-CFTR: flexibility, with frequent transient formation of interdomain hydrogen bonds, and loosely stacked aromatic sidechains, (F1068, R1070W, and F1074, mimicking F1068, F508 and F1074 in wild-type CFTR). F508del-CFTR had a distorted aromatic stack, with F1068 displaced towards space vacated by F508. In F508del/R1070F-CFTR, which largely retained F508del defects, R1070F could not form hydrogen bonds, and the interface was less flexible. Other ICL4 second-site mutations which partially rescued F508del-CFTR are F1068M and F1074M. Methionine side chains allow hydrophobic interactions without the steric rigidity of aromatic rings, possibly conferring flexibility to accommodate the absence of F508 and retain a dynamic interface. Finally, two mutations identified in a yeast scan (A141S and R1097T, on adjacent transmembrane helices linked to ICL1 and ICL4) also partially rescued F508del-CFTR function. These studies highlight the importance of hydrophobic interactions and conformational flexibility at the ICL4/NBD1 interface, advancing understanding of the structural underpinning of F508del dysfunction.

## Introduction

The ATP-binding cassette (ABC) transporter family is a large superfamily of proteins (1, 2). Within this superfamily, the cystic fibrosis transmembrane conductance regulator, CFTR (3), is the only protein known to function as an ion channel (4). Nevertheless, CFTR shares structural features with other ABC transporters. In particular, CFTR’s two asymmetric nucleotide-binding domains, NBD1 and NBD2, interact with two transmembrane domains (TMD1 and TMD2) possessing an almost typical Type IV fold (5). Interactions between the TMDs and NBDs are mediated by two pairs of intracellular loops (ICLs). The units formed by a surface depression in each NBD and the two ICLs that contact it, have been described as ‘ball-and-socket joints’ (6). While NBD1 forms a relatively shallow socket that interfaces with ICL1 and ICL4, NBD2 forms a deeper socket which contacts ICL2 and ICL3. ICL2 and ICL4 cross over to the NBD linked to the opposite TMD, forming a domain-swapped arrangement like that found in other Type IV ABC transporters, such as bacterial Sav1866 (7–9), MsbA (10), TM287/288 (11), mammalian P-glycoprotein (12), human ABCB10 (13), and McjD (14). In CFTR, interactions at the NBD-TMD interfaces are responsible for coupling ATP binding and hydrolysis to channel gating by transmission of conformational changes from the NBDs to the TMDs, which form the anion-selective permeation pathway (4).

CFTR plays an important physiological role, controlling epithelial secretions in several organs, such as the airways, intestine, pancreas, biliary ducts, sweat glands and reproductive tracts (15). Cystic fibrosis (CF), caused by loss of function mutations in the *CFTR* gene, is the most common life-limiting genetic disease in populations of European descent, affecting 1 in ∼2500-3000 newborns (16–18). CF-causing mutations are unequally distributed between the two NBD-TMD interfaces (6): 16 are found at the TMD/NBD1 interface, but only 5 at the TMD/NBD2 interface (list of CFTR2 variants 31 July 2020, https://cftr2.org/mutations_history). Possibly the absence of a short helix in NBD1 – present in NBD2 and in the NBDs of other ABC transporters (19) – is responsible for a shallower socket and weaker ICL4/NBD1 interactions, rendering the ICL4/NBD1 interface particularly vulnerable to harmful mutations. Moreover, a more dynamic NBD1 structure around the socket might also contribute to causing this mutation hotspot (20).

By far the most common CF-causing variant is F508del, which deletes a phenylalanine in NBD1 that contributes molecular contacts at the ICL4/NBD1 interface (16, 21, 22). Even though X-ray structures of human F508del NBD1 show that there are only small local changes in the conformation of the loop comprising residues 507 to 511 (23), the deletion has great impact on the biogenesis and function of CFTR. While wild-type (WT) CFTR becomes complex-glycosylated in the Golgi, the F508del mutant does not undergo any detectable complex glycosylation at 37 °C (24–30). Instead, misfolded F508del-CFTR is trapped in the endoplasmic reticulum (ER), ubiquitinated and then degraded by the proteasome (31).

The minute amount of F508del-CFTR that escapes to the plasma membrane has decreased membrane stability (32–34) and a severe gating impairment. The latter is characterized by a reduction in open probability (*P*_o_) (30, 35–40) caused by a prolonged closed time interval between bursts. The F508del mutation does not strongly affect CFTR pore properties such as single-channel conductance, and anion selectivity, although a reduction in current amplitude can occur once instability develops (reviewed in (41)).

In the laboratory, a common strategy to promote F508del-CFTR trafficking to the plasma membrane involves incubating cells at low temperature, 26–30 °C (30, 34, 42–44). Low temperature provides an energetically favourable F508del-CFTR folding (30) and proteostasis (43) environment, leading to a decrease in misfolding and improved trafficking to the plasma membrane. Moreover, evidence suggests that low temperature promotes trafficking of immature (core-glycosylated) F508del-CFTR, via a nonconventional trafficking pathway that by-passes the Golgi (44, 45).

Another laboratory strategy for F508del-CFTR rescue, is to introduce second-site (revertant) mutations in *cis* with F508del. Identified revertant mutations located in NBD1 include: V510D/E/A (46–48), I539T (49, 50), G550E (49), R553M/Q (51) and R555K (50, 52). The latter mutations reduce F508del-NBD1 thermodynamic and kinetic instability (53, 54). By contrast, R1070W, located in ICL4 (53, 55, 56), acts by restoring interactions at the ICL4/NBD1 interface (55, 57).

Here, we investigated F508del-CFTR rescue by second-site mutations. We systematically scanned positions 1064-1074 of ICL4, substituting native amino acids with F, H, M, Q, W and Y. In addition, we tested the effects of A141S and R1097T. These mutations correspond to the revertant mutations F270S (58) and R1116T (59) identified in F670del-Yor1p, a yeast homolog of F508del-CFTR in which the deletion of F670 causes defects similar to those of F508del in CFTR (58). The panel of mutants was studied in live HEK-293 cells using a new high-content assay (60), allowing simultaneous quantification of CFTR cellular conductance and the amount of CFTR in close proximity to the plasma membrane. Our screen validates R1070W as a particularly effective F508del-CFTR revertant.

To investigate how the tryptophan substitution improves biogenesis and ion-channel function so effectively, molecular dynamics simulations were run using systems representing the F508del-CFTR mutant, the F508del/R1070W-CFTR revertant, and the much less effective F508del/R1070F-CFTR revertant. The simulations were compared to those obtained with a WT CFTR system (61). Our results reveal how the R1070W revertant mutation might restore transient interactions between ICL4 and the F508del-NBD1 loop, including a network of hydrophobic interactions between aromatic residues at the ICL4/NBD1 interface, as seen in WT CFTR. Our mutagenesis scan results together with the altered molecular dynamics at the ICL4/NBD1 interface *in silico*, suggest hypotheses to explain the molecular basis of the F508del-CFTR defects and how they might be repaired.

## Results

To assess how second-site mutations in *cis* with F508del affect the CFTR anion channel, YFP(H148Q/I152L)-CFTR fluorescence quenching in response to extracellular I^-^ addition was quantified following expression in HEK-293 cells. A 24 hour incubation at 28 °C was included, to minimize misfolding and optimize trafficking, and the low temperature was maintained throughout image acquisition. Whole cell conductance (G) and membrane potential (V_m_) at steady-state, immediately preceding I^-^ addition, were estimated by fitting of a mathematical model to the quenching time course. Moreover, we simultaneously measured CFTR membrane proximity by quantifying the YFP(H148Q/I152L)-CFTR fluorescence located at the border of cells, using mCherry fluorescence as an internal standard for comparison (60). **Table S1** summarizes the assay readouts.

### Whole-cell conductance (G)

In the control (DMSO) condition, mutants with the F508del background typically had an average G estimate of around 1 nS (M = 0.97 nS, SD = 1.29, N = 64), consistent with a small anion permeability reflecting endogenous, non-CFTR-mediated conductances and minimal basal phosphorylation of CFTR. Following CFTR activation (through the cAMP pathway) by 10 µM forskolin (**Figure 1A**), the anion conductance of WT CFTR increased significantly (forskolin: *Mdn* = 117.70 nS, *N* = 17; DMSO: *Mdn* = 2.35 nS, *N* = 20; see one-tailed Wilcoxon ranking tests in **Table S2**). For F508del-CFTR too, there was a modest, albeit significant, increase in G after addition of forskolin (*Mdn* = 5.7 nS, *N =* 18) compared to the control condition (*Mdn* = 0.86 nS, *N =* 19). An increased conductance after addition of forskolin compared to DMSO was observed in 24 of 61 mutants with second-site mutations in *cis* with F508del at the ICL4/NBD1 interface (**Table S2, Figure S3**).

**Figure 1.**
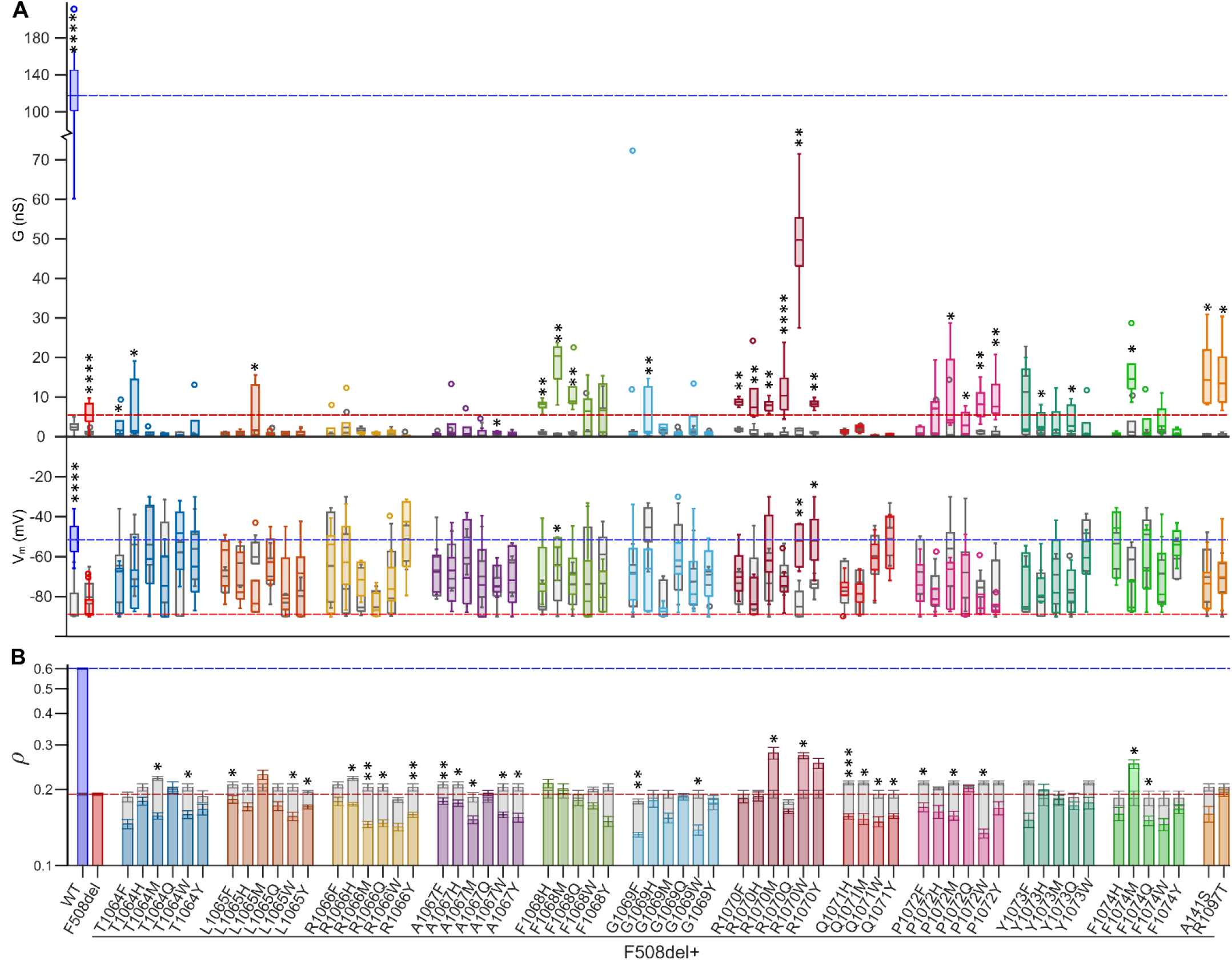
Effects of second-site mutations in *cis* with F508del on CFTR conductance, membrane potential, and CFTR membrane proximity. **A**) CFTR conductance (G in nS, upper panel) and membrane potential (V_m_ in mV, lower panel), measured in HEK-293 cells expressing WT CFTR, F508del-CFTR, or F508del-CFTR with second-site mutations. The dashed reference lines indicate the median conductance (upper panel) and membrane potential (lower panel) after steady-state activation with forskolin of WT CFTR (blue) and F508del-CFTR (red). Asterisks show results of one-tailed Wilcoxon ranking tests used to compare G and V_m_ after addition of DMSO (control; gray boxes) vs. 10 µM forskolin (colored boxes). **B**) Measurements of CFTR membrane proximity (ρ) obtained from the same CFTR-expressing HEK-293 cells described in **A**. Dashed reference lines indicate the average CFTR membrane proximity for cells expressing WT CFTR (blue) and F508del-CFTR (red). Mean log_10_ρ values were paired per plate and paired t-tests were performed on the mean log_10_ρ measurements of cells expressing F508del-CFTR with (colored bars) vs. without (gray bars) second-site mutation.

**Table S4** demonstrates that many second-site mutations further impaired F508del-CFTR function – especially substitutions at sites T1064, L1065, R1066, A1067, G1069 and Q1071. By contrast, eight mutations significantly increased the F508del-CFTR-mediated, forskolin-stimulated conductance (Wilcoxon Rank Sum tests, **Table S4**). Among these, R1070W, was particularly effective, increasing F508del-CFTR conductance to 42% of the value measured for WT CFTR (*Mdn* = 49.76 nS, *N =* 5). As a comparison, chronic treatment of F508del-CFTR expressing HEK-293 cells with the clinically-approved CFTR corrector lumacaftor (3 μM VX-809 for 24 h at 28 °C) resulted in a whole-cell conductance of only 12% of that measured for WT CFTR (*Mdn* = 13.73 nS, *N =* 7, data not shown). After R1070W, the most successful revertant mutations were F1068M and F1074M, followed by A141S and R1097T. Finally, R1070Q, F1068Q and R1070F mutations all gave smaller, but still significant, improvements in G (**Table S4**).

### Membrane potential (V_m_)

More than half of the HEK-293 cells expressing second-site mutations resulted in a significantly depolarized V_m_ after steady-state activation with 10 µM forskolin when compared to those expressing F508del-CFTR (Wilcoxon Rank Sum tests, **Table S4**). However, for most of these cells, a relatively depolarized V_m_ was also present under control conditions. A significant depolarisation of V_m_ resulting from activation of CFTR with forskolin was seen only for WT CFTR, F508del/R1070W, F508del/R1070Y, and F508del/F1068M (one-tailed Wilcoxon ranking tests (**Table S2**).

### Membrane proximity (ρ)

Watershed-based image segmentation on the mCherry images allowed us to approximate the location of the plasma membrane without relying on efficient YFP(H148Q/I152L)-CFTR membrane trafficking. For each cell, the amount of CFTR in proximity to the membrane, denoted as ρ, was defined as the ratio of the average normalized YFP(H148Q/I152L)-CFTR fluorescence intensity within the membrane proximal zone (a ∼ 1 µm wide band adjacent to the cell boundary) to the average normalized mCherry fluorescence intensity throughout the entire cell (ρ = f_YFP membrane_/f_mCherry cell_). This metric informs about first, trafficking (the fraction of YFP(H148Q/I152L)-CFTR reaching the membrane, f_YFP membrane_/f_YFP, cell_) and second, the fusion protein’s metabolic stability with respect to mCherry’s (quantified by the ratio f_YFP cell_/f_mCherry cell_), a function of overall rates of biosynthesis and degradation. The ρ measurements approximated a lognormal distribution and were log_10_ transformed before determining plate means for each mutation. To evaluate the effects of second-site mutations on F508del-CFTR delivery to the plasma membrane, we compared the mean log_10_ρ measurements of cells expressing F508del-CFTR in the absence and presence of second-site mutations (**Figure 1B, Table S5**). Only R1070W, R1070M and F1074M significantly increased F508del-CFTR membrane proximity, whereas it was significantly reduced by 25 substitutions (**Figure 1B, Table S4**).

### Gating and conduction properties

CFTR conductance increases approximately linearly with membrane proximity (60), consistent with the ρ metric being proportional to the number of channels at the plasma membrane. Because whole-cell conductance (G) is the product of the number of channels at the membrane (*N*), open probability (*P*_o_) and single-channel conductance (*γ*), our two assay readouts allow evaluation of the gating and conduction properties (*P*_o_·*γ*) of CFTR channels located at the plasma membrane (see (60)). The dotted lines in **Figure 2** describe conductance as a function of membrane proximity assuming single-channel properties (*P*_o_·*γ*) of WT CFTR after steady-state activation with forskolin, with the specific experimental conditions used (see **Figure S6**). Data points falling above or below the line are suggestive of single-channel activity higher or lower, respectively, than those of WT CFTR.

**Figure 2.**
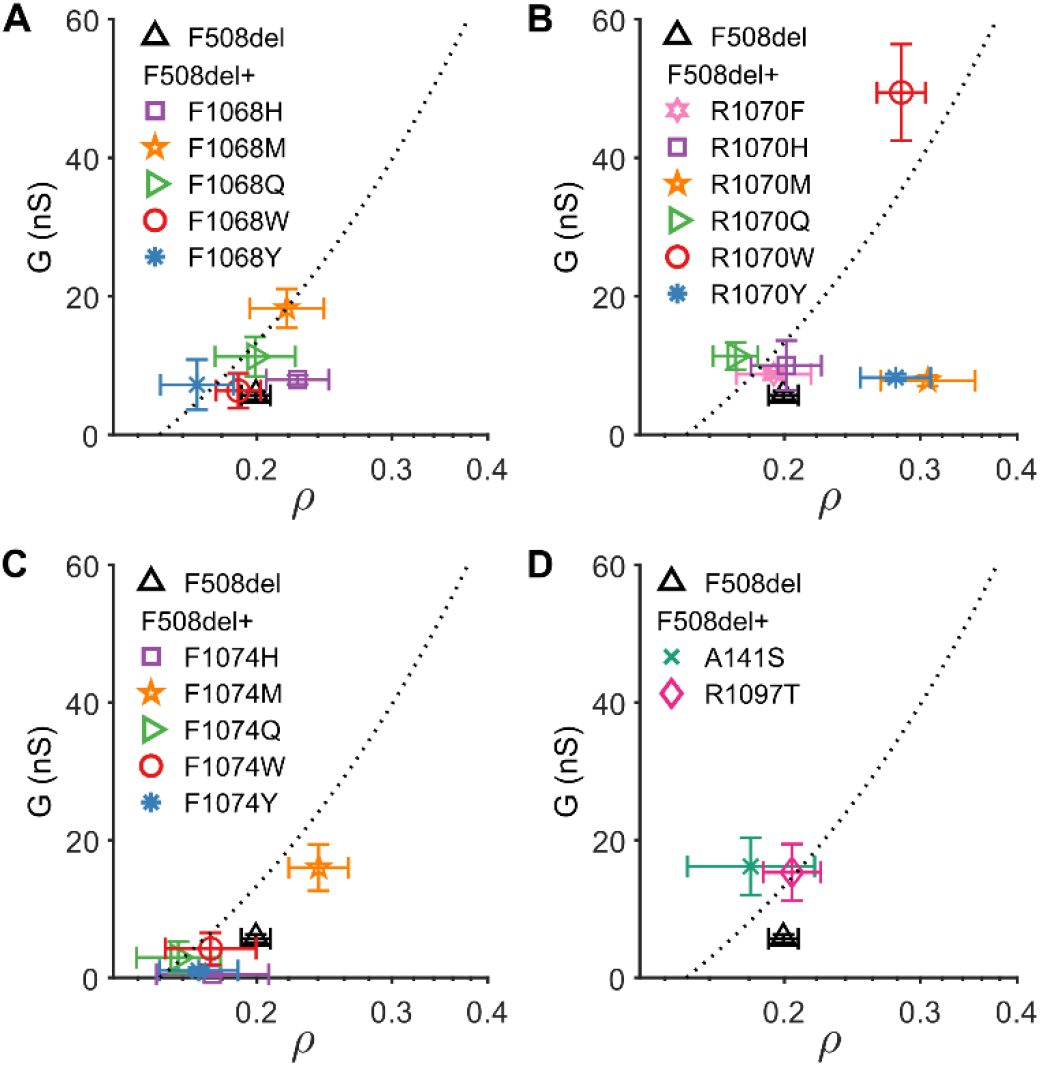
Plate matched G-ρ Measurements. G was plotted as a function of membrane proximity (ρ, obtained by back transformation of mean log_10_ρ). The panels show G-ρ plots for F508del-CFTR in the absence and presence of second site mutations at residues F1068 (**A**), R1070 (**B**), F1074 (**C**), and identified in the Yor1p screen (A141S and R1097T, **D**). Markers and error bars represent the mean G ± S.E.M. (y-axis) and back transformed mean ρ ± upper and lower limits of the S.E.M. (x-axis).

F1068M (orange five-point star, **Figure 2A**) improves the whole-cell conductance of F508del-CFTR. The rightward (non-significant) shift on the ρ-axis positions the point on the regression line, suggesting that *P*_o_·*γ* characteristics of F508del/F1068M-CFTR might be close to those of WT CFTR. F508del/R1070M (orange five-point star, **Figure 2B**) is significantly shifted to the right compared to F508del-CFTR, suggestive of improved biogenesis, trafficking and/or membrane stability. However, the mutant falls far below the regression line, suggesting that even though the number of channels at the plasma membrane has increased *P*_o_·*γ* is still much lower than that of WT CFTR. By contrast, F508del/R1070W (red circle, **Figure 2B**) falls above the regression line, consistent with published single-channel recordings demonstrating rescue of *P*_o_ to WT values (while *γ* is unchanged in both F508del-CFTR and F508del/R1070W, (62). All substitutions at site F1074, except for methionine (M; orange 5-point star, **Figure 2C**), decreased conductance, such that the forskolin-dependent increase in G after activation seen in F508del-CFTR is lost (**Table S2**). By contrast, the F1074M substitution not only significantly increased conductance, but was also one of the three mutants that significantly increased the membrane proximity of F508del-CFTR. Part of the increase in whole-cell conductance is thus due to an increase in the number of channels at the plasma membrane. F508del/A141S (green cross) and F508del/R1097T (pink diamond) (**Figure 2D**) both increase conductance relative to F508del-CFTR. However, neither mutation improves membrane proximity, suggesting a G increase dependent on improved channel function.

### Molecular dynamics simulations

We used molecular dynamics (MD) simulations to investigate whether efficacy of rescue was correlated with restoration of the interactions between ICL4 and NBD1 found in WT CFTR. For comparison to WT CFTR, we used previously published simulations of the WT zebrafish CFTR (zCFTR) (61), whose sequences at the ICL4/NBD1 interface differ from those of hCFTR at only six positions in the NBD1 loop including F508 (positions 495-512, human numbering) and eight in ICL4 (positions 1050-1080, **Table S7**). In subsequent sections, we use the numbering of residues in hCFTR to refer to the positions of residues in both hCFTR and zCFTR (e.g. zR1070 indicates the zCFTR residue corresponding to R1070 in hCFTR, i.e. R1078, see **Table S7, Figure S8**). We selected for analysis two ICL4 second-site mutations, which rescue F508del-CFTR with different efficacy: R1070W that improved markedly both biogenesis and conductance, and R1070F, which has only minor effects on conductance.

We constructed a model for the F508del-CFTR system by replacing NBD1 in the ATP-bound structure of hCFTR (63) with the experimental structure of F508del-NBD1 (64). The resulting system is hereafter termed F508del/R1070. We then generated two additional F508del systems by replacing R1070 with (i) a tryptophan residue (F508del/R1070W), and (ii) a phenylalanine residue (F508del/R1070F). Each system was embedded in a POPC lipid bilayer and simulated for 2 μs. The ICL4/F508del-NBD1 interfaces were compared with the ICL2/NBD2 interface of the same systems, as well as with previously published, ATP-bound and ATP-free simulations of the WT zCFTR (6, 61, 65).

In our MD simulations, we assume that the conformations sampled by the mutant proteins, which reach the plasma membrane are not grossly different from those adopted by WT CFTR. Although mutation effects during biogenesis might result in proteins that fold differently, these effects are minimized by low temperature incubation. Moreover, the significant increase in conductance following stimulation by forskolin (**Table S2**) for these three mutants suggests that a large proportion of the mutant channels at the plasma membrane retain regulation of gating by cAMP-dependent phosphorylation.

#### RMSD-based cluster analysis

First, we investigated whether the presence of aromatic residues at the R1070 position alters the structure of the F508del-NBD1 loop. To address this aim, we analysed the conformation of the F508del-NBD1 loop by means of a RMSD-based cluster analysis of the ICL4/NBD1 interface (ICL4 residues 1050-1080 and F508del-NBD1 loop residues 495-512). With a RMSD cut-off of 0.1 nm, we detected: (i) 39 clusters for the F508del/R1070 system, with the first cluster and the top 3 clusters (**Figure 3A-C**) representing approximately 43% and 69%, respectively, of the total structures; (ii) 35 clusters for the F508del/R1070W system, with the first cluster and top 3 clusters (**Figure 3D-F**) representing approximately 48% and 73%, respectively, of the total structures; (iii) 9 clusters for the F508del/R1070F system, with the first cluster alone representing approximately 89% of the total structures (**Figure 3G**). The same analysis was performed on datasets from simulations on WT zCFTR (**Figure 3H-I**), with cluster 1 representing 93% and 74% of the total structures from the ATP-bound and ATP-free simulations, respectively. **Figure 3** demonstrates that the NBD1 loop in the F508del/R1070 and F508del/R1070W mutants explored more extended conformations following the helical part (residues 502-507) of the loop. This can be seen for cluster 1 of the F508del/R1070 system (**Figure 3A**) and for the first two clusters of the F508del/R1070W system (**Figure 3D-E**). More compact conformations, similar to the WT hCFTR NBD1 loop conformation (**Figure S8**), were retrieved for (i) clusters 2 and 3 of the F508del/R1070 system (**Figure 3B-C**); (ii) cluster 3 of the F508del/R1070W system (**Figure 3F**), and (iii) the F508del/R1070F system (**Figure 3G**). These folded conformations were also retrieved in the most populated cluster of the ATP-bound and ATP-free zCFTR systems (**Figure 3H-I**).

**Figure 3.**
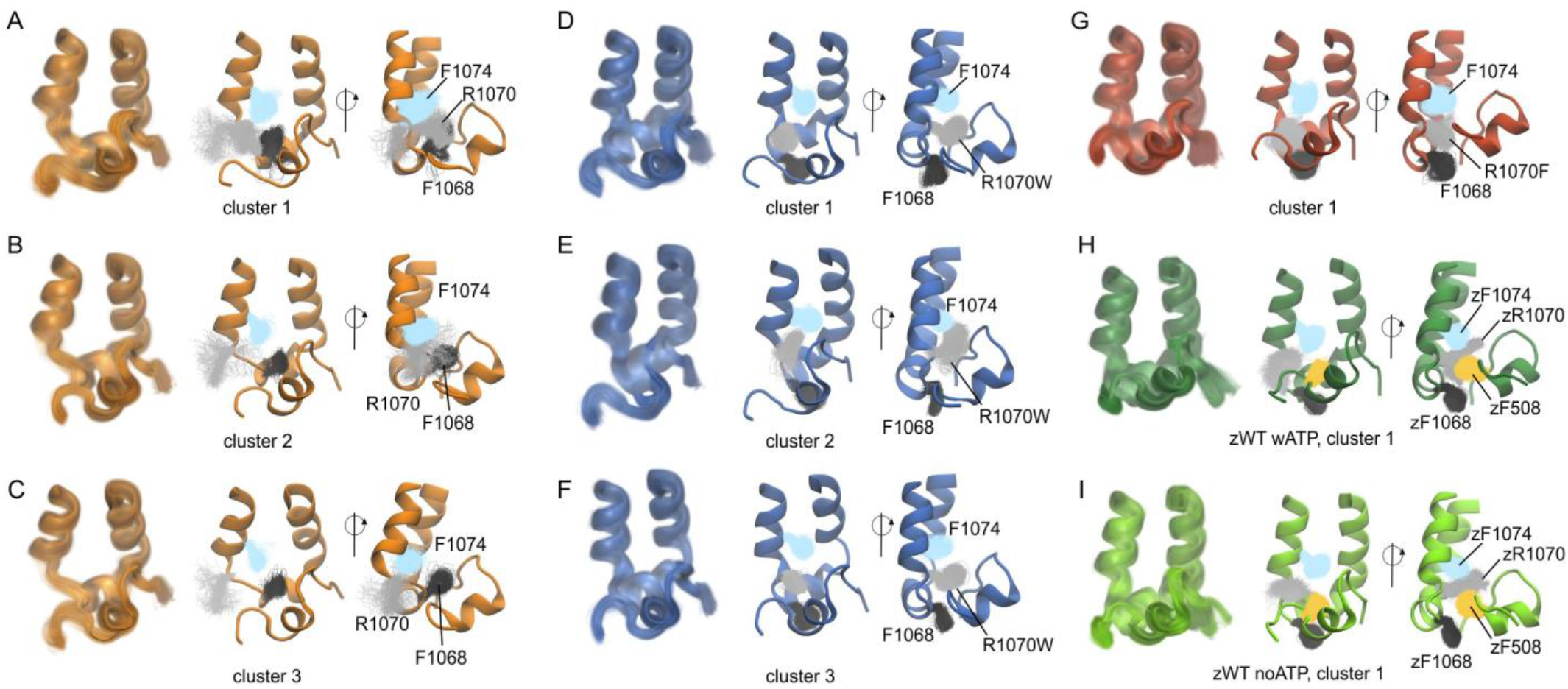
Dynamics of the ICL4/F508del-NBD1 interface. Results of the cluster analysis for the (**A**-**C**) F508del/R1070 simulation system; (**D**-**F**) the F508del/R1070W system; (**G**) the F508del/R1070F system; (**H**) ATP-bound and (**I**) ATP-free zCFTR systems previously simulated (61). For the F508del/R1070 and F508del/R1070W systems, the three most populated clusters are shown, while for F508del/R1070F and WT zCFTR only the first cluster is shown. For each cluster, the left panel is the front view of the interface, with the members of the cluster shown in transparent cartoons. The middle and right panels show the cluster centre as cartoon and the side chains of F1068 (black), R1070X (gray), F1074 (light blue), and F508 (yellow) of all the members of the cluster, as lines.

#### Aromatic residue side chain orientations

The interface between ICL4 and the NBD1 loop near R1070 is characterized by the presence of several aromatic residues, including F1068 and F1074 from ICL4 and F508 from NBD1. **Figure 3A-G** shows the side chain orientation of F1068 (dark gray), R1070X (light gray) and F1074 (light blue) of all the members of a given cluster. We considered the orientation of these interfacial residues by measuring the first rotamer (χ1) of their side chains over the entire simulation time (**Figure 4A, 4C**). χ1 for F1068 shows a peak near 300°, corresponding to the side-chain orientation of the most populated clusters for F508del/R1070F, F508del/R1070W, and WT zCFTR, corresponding to the “downwards” side chain orientation shown in **Figure 3D-I**. By contrast, for the F508del/R1070 system, the main peak is at 180°, reflecting the “upwards” movement of the side chain towards the NBD1 loop, as for the most populated clusters in **Figure 3A-C**. For the aromatic residues at the 1070 position, R1070W shows a bimodal distribution (middle panel, **Figure 4A)**, with the major peak near 280-300° and a smaller one at approximately 200-210°. The major peak corresponds to the side chain orientation shown in **Figure 3D, 3F**, and is shared with R1070F (**Figure 3G**), while the smaller peak corresponds primarily to structures that form the second most populated cluster of F508del/R1070W, with the tryptophan side chain flipped upwards (**Figure 3E**). No significant differences were retrieved across the three F508del systems for the χ1 distribution of F1074. At the ICL2/NBD2 interface, Y275 corresponds to the F1068 position in ICL4 (**Table S7**), and its χ1 distribution is centered near 300° (**Figure 4B, 4D**), similar to F1068 in F508del/R1070W and F508del/R1070F, and WT zCFTR. W277 at the ICL2/NBD2 interface corresponds to R1070 in ICL4 (**Table S7**), and its χ1 distribution shows a peak near 300° in all simulation systems, as for R1070F and the major peak of R1070W.

**Figure 4.**
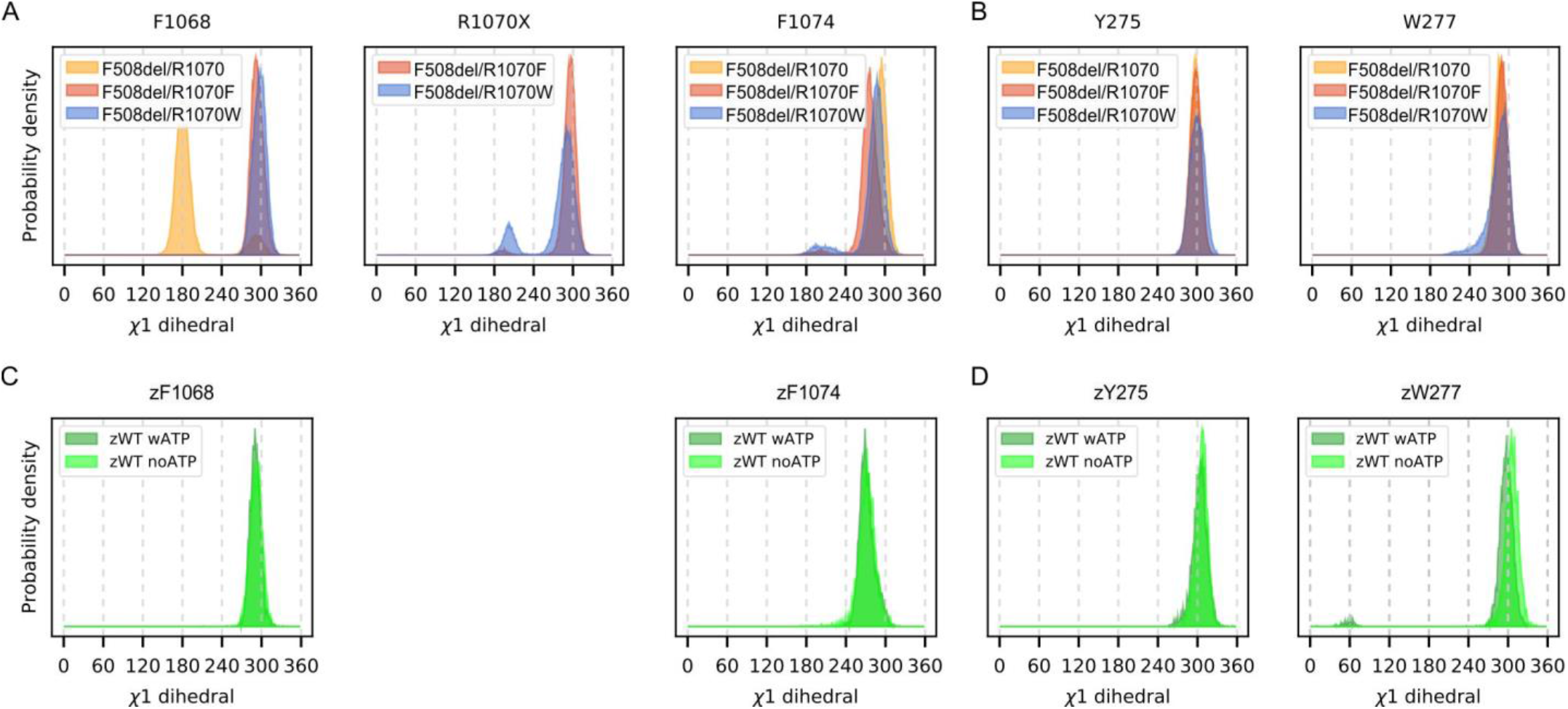
Aromatic side-chain orientations at TMD/NBD interfaces. Probability density of the χ1 dihedral angle for selected aromatic residues. (**A**) F1068, R1070X and F1074 from ICL4 in the F508del simulation systems; (**B**) Y275 and W277 from ICL2 in the F508del simulation systems; (**C**) zF1068 and zF1074 (equivalent to F1076 and F1082, zebrafish numbering) from ICL4 in the WT zCFTR simulation systems; (**D**) zY275 and zW277 (Y276 and W278) from ICL2 in the WT zCFTR simulation systems (61).

In summary, with the exception of F1068 in F508del/R1070, the χ1 angle of the selected residues at the interface corresponds to the distributions obtained from previous simulations (61) on the WT zCFTR structure (**Figure 4A, 4B** vs. **Figure 4C, 4D**).

#### Spatial overlap between F508 in WT CFTR and side chains at position 1070

To test for overlap of R1070F and R1070W with the region occupied by F508 in WT hCFTR, the ICL4/NBD1 interface from the hCFTR structure was superimposed on the center of cluster 1 for the F508del/R1070F system (**Figure S9A**) and those of the three most populated clusters of the F508del/R1070W system (**Figure S9B-D**). While R1070F projected towards the space vacated by F508 (**Figure S9A**), the overlap was greater for the F508del/R1070W system, particularly cluster 1 (**Figure S9B**). We also compared the orientation of R1070F and R1070W with that of W277 at the ICL2/NBD2 interface (**Figure S9E-H**). Similar results were obtained with this comparison; a higher degree of overlap was achieved with the R1070W mutant (**Figure S9F** vs. **Figure S9E**).

#### Hydrogen-bond interactions at the ICL4/F508del-NBD1 interface

In our previous WT zCFTR simulations (61), we observed that zR1070 was highly dynamic, oriented both towards and away from the interface, and forming transient interactions with zF508 and other residues of the NBD1 loop (**Figure 5A** and **Table S7**). We therefore investigated whether equivalent interactions were present in the F508del systems. We found that R1070 and R1070W formed hydrogen bonds with residues E504, I507 and G509 of the NBD1 loop. Interactions also occurred between Y1073 and E504 (**Figure 5B**). Overall, hydrogen-bond interactions in the F508del systems, while not lasting for the entire simulation time, were more frequent and persistent than those in the WT zCFTR system.

**Figure 5.**
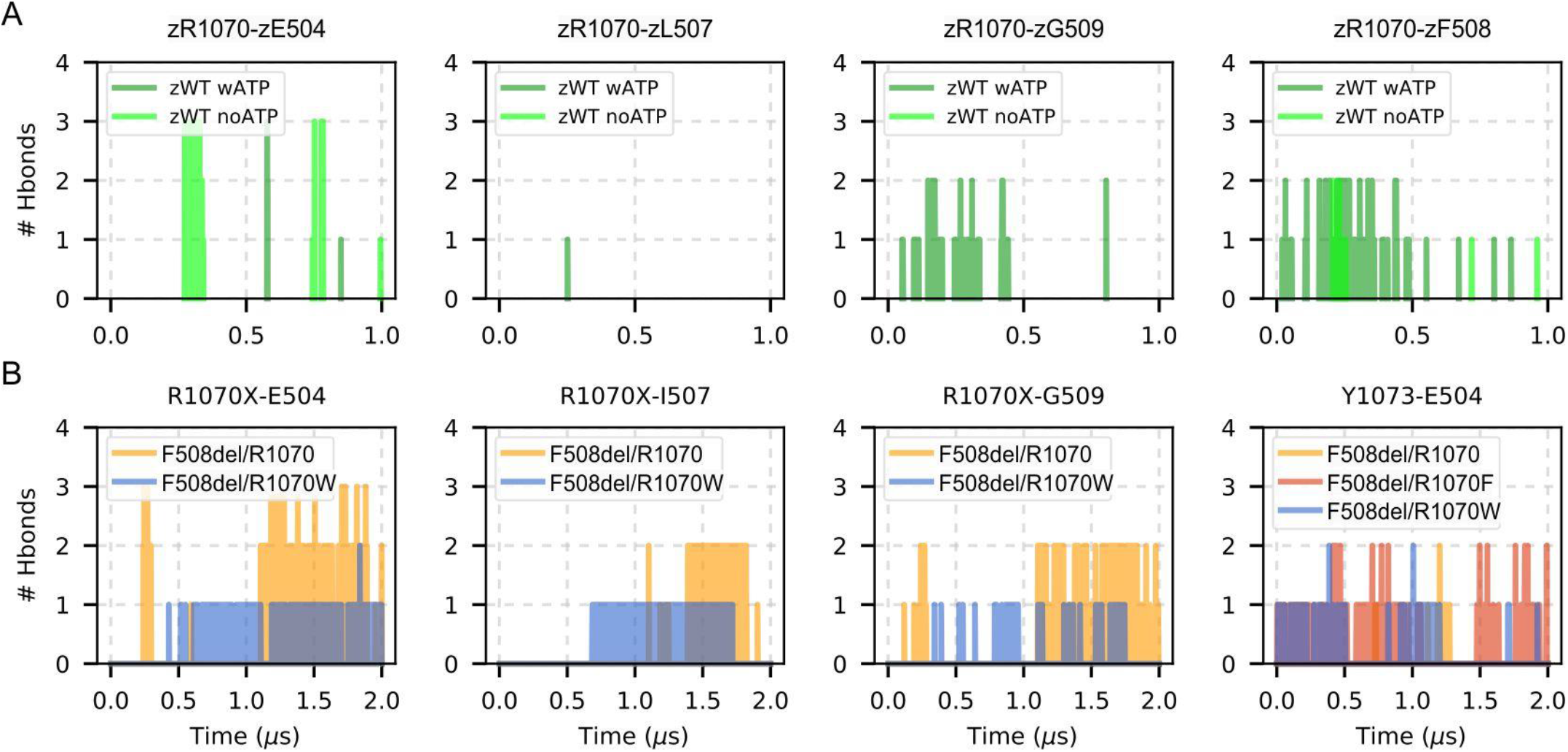
Hydrogen bonds between ICL4 and NBD1 residues. Hydrogen bond interactions were detected as a function of time for selected pairs of residues at the ICL4/NBD1 interface. (**A**) Interactions at the ICL4/NBD1 interface in the WT zCFTR simulations (61), zWT wATP and zWT noATP indicate the WT zCFTR simulation of the ATP-bound and ATP-free structures, respectively. (**B**) Interactions detected at the ICL4/F508del-NBD1 interface in the three F508del systems studied.

## Discussion

Our systematic, empirical scan of ICL4, seeking mutations capable of repairing F508del defects, highlighted a number of second-site revertant mutations. The replacement of R1070 with a tryptophan (53, 55, 56) was found to be most effective at increasing plasma membrane levels and function of F508del-CFTR. This confirms patch-clamp recordings, which show how the R1070W mutation restores F508del-CFTR plasma membrane stability and gating kinetics to levels measured for WT CFTR (62). Using molecular dynamics simulations, the structural basis of this revertant mutation’s action was investigated. Our systematic scan also identified a small number of new revertant mutations, which mitigated the defects of F508del to some degree. Taken together, our results underscore the importance, for CFTR biogenesis and function, of a dynamic ICL4/NBD1 interface characterized by multiple transient hydrophobic and hydrogen-bond contacts.

### F670del-Yor1p revertants

We introduced CFTR equivalents of the F670del-Yor1p revertant mutations F270S (66) and R1116T (59) into F508del-CFTR. In hCFTR, the corresponding revertant mutations are A141S (in transmembrane helix 2, TM2) and R1097T (in TM11), respectively. Both revertant mutations, situated on adjacent TMs at the same horizontal plane within the membrane-embedded portion of the protein, significantly rescued F508del-CFTR activity. Structurally, the two residues are coupled to the TMD/NBD1 ball-and-socket joint, through helical portions of TM2 (linking to ICL1) and TM11 (linking to ICL4) (67). F270S only rescued the folding and trafficking of F670del-Yor1p in combination with another revertant mutation, R1168M, in TM12 (66). By contrast, A141S, by itself, significantly rescued the conductance of F508del-CFTR, albeit, it was without effect on its membrane proximity, suggesting an effect on *P*_o_ and/or *γ*. Unlike F270S, R1116T, by itself, increased F670del-Yor1p function. Our data show that R1097T too increased F508del-CFTR conductance by improving its gating and/or conduction properties. While we cannot completely rule out an effect on conduction, residues in TM2 and TM11 do not play a major role in lining the inner vestibule of the pore (68) (but note (69)), suggesting effects on *P*_o_ are more likely. It is possible that in F670del-Yor1p, as in F508del-CFTR, the enzymatic cycle time is greatly prolonged by a slow transition from the inward-facing to outward-facing conformation (corresponding to opening in CFTR, (70–72)). R1116 and R1097 are linked, via TM11, to the TMD/NBD1 interface altered by the phenylalanine deletion. The threonine substitutions might facilitate coupling of NBD dimerisation to the TMD rearrangement resulting in an outward-facing conformation. Further studies are required to better understand the mechanistic details of this rescue.

### Why is R1070W such an effective revertant mutation?

Introducing the R1070W mutation into the F508del-CFTR background greatly increased the CFTR-mediated anion conductance. In an attempt to identify the structural basis of this remarkably efficient rescue, we compared three different MD simulations of F508del systems (F508del/R1070, F508del/R1070W and F508del/R1070F) with simulations previously performed on the WT zCFTR systems (61). While other MD simulations have analysed the NBD dimer interface (73), we focused on the ICL4/NBD1 interface. Two potentially important features of protein dynamics emerged, suggesting that both the indole nitrogen and the aromatic ring in R1070W play important roles (**Figure 6**).

**Figure 6.**
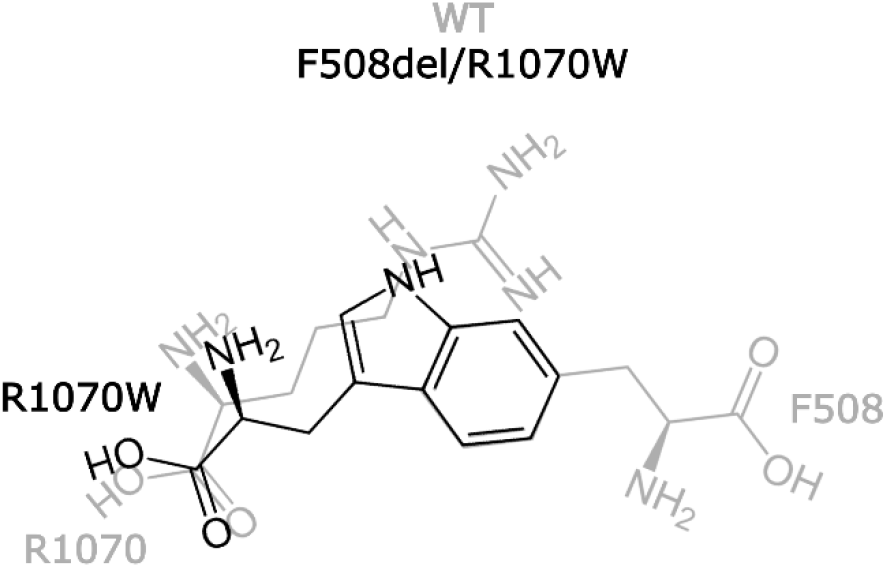
Schematic overview of the R1070W mutation introduced in *cis* with F508del. In the F508del/R1070W mutant the tryptophan at site 1070 might provide appropriately positioned (i) hydrogen-bond donor (the indole nitrogen atom) and (ii) aromatic ring, restoring hydrophylic and hydrophobic contacts lost by deletion of F508 and replacement of R1070 (F508 and R1070 in WT CFTR shown in light gray).

#### F508del/R1070W restores ICL4/NBD1 aromatic interactions

In WT CFTR, the interface between ICL4 and NBD1 is characterized by the presence of several aromatic residues. In MD simulations of WT zCFTR, we found that the side chains of zF1068, zF508, and zF1074 loosely stack in an alternating fashion (**Figure 3H, 3I**). In the MD simulation of F508del/R1070, the arrangement of aromatic side chains at the ICL4/NBD1 interface deviated noticeably from this pattern, due to the upward movement of F1068 towards the region vacated by the deletion of F508. Our MD simulations of F508del/R1070W and F508del/R1070F showed that both R1070W and R1070F side chains occupy the region of space sampled by F508 in WT zCFTR (**Figure 3D-G**). As a result, the loose stacking of aromatic residues in WT zCFTR and hCFTR is restored. The bulkier side chain of R1070W permitted better overlap with the region occupied by F508, particularly for the most populated cluster of F508del/R1070W, comprising 48% of the total structures (**Figure S9**). It is interesting to note that at the homologous ICL2/NBD2 interface, a tryptophan, W277 (ICL2), corresponds to R1070, and projects its side chain towards the NBD2 loop (positions 1303-1309), where a proline (P1306) is present at a position equivalent to F508. A tryptophan-proline pair is completely conserved at these positions at the ICL2/NBD2 interface among a set of asymmetric ABC transporter sequences (74). Similarly, an aromatic side-chain is complet ely conserved at a position equivalent to F508 at the ICL4/NBD1 interface. However, a basic side chain at position equivalent to R1070 is much less conserved (74).

#### F508del/R1070W restores ICL4/NBD1 hydrogen-bond interactions, allowing flexibility of the NBD1 loop

In WT zCFTR, the NBD1 loop mostly adopts a compact, folded structure, but several residues in ICL4, including zR1070, are highly dynamic and form transient hydrogen bonds with residues in the NBD1 loop, including zF508 (**Figure 5A**, see also (22, 75, 76)). Our MD simulations of the three F508del systems highlighted a clear dichotomy. F508del/R1070 and F508del/R1070W had a more flexible NBD1 loop, sampling more extended conformations, and forming transient hydrogen bonds with residues of ICL4. By contrast, F508del/R1070F adopted a more rigid interface, with no groups at the 1070 position capable of establishing ICL4/NBD1 hydrogen bonds (**Figure 5B**). R1070W orientated towards the interface occupying the region sampled by zR1070 in WT zCFTR, formed similar hydrogen bond contacts, albeit more frequently (**Figure 5A** vs. **5B**). R1070F adopted a similar orientation. However, because hydrophilic contacts are not possible for the phenylalanine side-chain, the ICL4/NBD1 interface in this mutant is fixed and cannot sample alternative conformations with similar stability. This is demonstrated by the absence of significant changes in the NBD1-loop conformation throughout the simulation (the top cluster, **Figure 3G**, includes 89% of all structures). Functionally, a dynamic ICL4/NBD1 interface is crucial for CFTR gating, as demonstrated by the rapid and reversible interruption of channel activity that occurs upon formation of covalent cross-links between F508C and F1068C in a cys-less CFTR background (21). In addition, recent studies suggest that a frequent “uncoupling” of NBD1 is an integral feature of WT CFTR gating (20, 77). A tryptophan side-chain at position R1070, capable of both hydrogen bonding and hydrophobic contacts, would allow the protein to adopt conformations in which the interface is alternatively buried or more solvent exposed.

### Methionine substitutions at positions F1068 and F1074

Following R1070W, the most effective F508del-CFTR revertant mutations are F1068M and F1074M. These results are consistent with a crucial role played by the aromatic cluster at the core of the ICL4/NBD1 interface (including F1068, F508 and F1074) highlighted by our MD simulations and analysis (**Figure 3, Figure 4**). Methionine-aromatic interactions can occur at longer distances and are potentially more robust compared to purely hydrophobic contacts or salt bridges (78). The linear (non-branched) aliphatic chain that includes a sulphur atom gives more flexibility than any other hydrophobic side-chain (79), possibly allowing strong interactions with the aromatic cluster, despite the distortions caused by the absence of F508.

In addition, F1068M and F1074M might allow transient polar interactions (80, 81). Strikingly, one of the critical phenylalanines coordinating permeating anions in the Fluc-Ec2 channel can only be functionally replaced by methionine (82). We speculate that rescue by the introduced methionines in CFTR might rely on a dynamic switching between relatively strong hydrophobic interactions and interactions with polar residues of the NBD1 loop or with the solvent.

### Potential translational impact of this study

The results of the mutagenic scan presented here advance our understanding of the defect caused by absence of phenylalanine 508, the variant carried by most people with CF of European descent. As our understanding of the structure and molecular dynamics of the CFTR protein improves, information on the detailed impact of the deletion will inform efforts to design future therapies capable of optimizing the rescue of CFTR biogenesis and function.

The strong validation our data provides of the effectiveness of the revertant mutation R1070W might invigorate efforts to design novel gene therapy treatments for CF. Partial restoration of function might be achieved by cytosine base editing (transforming the arginine-encoding codon CGG to the tryptophan-encoding TGG). Base editing is more efficient and less prone to error than homology-directed repair ((83) and PT Harrison, personal communication). In addition, R1070W rescue of F508del-CFTR channels is likely to have synergistic effects with Class I and Class III correctors (57, 84) potentially allowing a simplification of modulator therapy. Thus, base-editing gene therapy might lead to benefits for F508del homozygous patients, in the *interim*, while the homology-directed repair gene therapy technology is advanced.

## Experimental Procedures

### Plasmid and site-directed mutagenesis

Mutations were introduced in the pIRES2-mCherry-YFPCFTR plasmid (see (60)) with help of complementary primers containing mutations (Eurofins MWG Operon, Germany) using the QuikChange protocol for site-directed mutagenesis (Stratagene). Sanger sequencing either outsourced to SourceBioscience (Nottingham, UK) or by the UCL Sequencing Facility with a 3100-Avant Genetic Analyzer (Applied Biosystems) was used to confirm the introduced mutations.

### Cell culture, transfection and incubations

HEK-293 cells were maintained at 37 °C in Dulbecco’s modified Eagle’s medium, supplemented with 2 mM L-glutamine, 100 units/mL penicillin and 100 μg/mL streptomycin, and 10% fetal bovine serum (all from Life Technologies, Inc.). Cells were seeded in black-walled 96-well plates (Costar, Fisher Scientific) coated with poly-D-lysine and transiently transfected with the pIRES2-mCherry-YFPCFTR plasmid using Lipofectamine 2000 (Life Technologies), following the manufacturer’s instructions. After transfection, cell plates were incubated at 37 °C for 24 h, then at 28 °C for a further 24 h to minimize misfolding (30). Before imaging, cells were washed twice in 100 μL of standard buffer (140 mM NaCl, 4.7 mM KCl, 1.2 mM MgCl_2_, 5 mM HEPES, 2.5 mM CaCl_2_, 1 mM glucose, pH 7.4).

### Image acquisition

The ImageXpress Micro XLS (Molecular Devices), an automated inverted wide-field fluorescence microscope with a temperature-controlled chamber (set to 28 °C), was used for image acquisition. Protocols for automated imaging and fluid additions, were created using MetaXpress software (Molecular Devices). A 20× objective was used to take 16-bit images of both mCherry (excitation/emission filters at 531 ± 20 / 592 ± 20 nm) and YFP-CFTR (excitation/emission filters at 472 ± 30 / 520 ± 35 nm). To evaluate CFTR activity at steady-state, images of mCherry and YFP fluorescence were taken every 2 s. After following the baseline for 20 s, CFTR was activated by the addition of 50 μl of standard buffer containing forskolin (10 μM final concentration) or DMSO (control, 0.05 %). After a further 230 s, when CFTR is expected to be gating at steady-state, 50 μL of I^-^ buffer (as standard buffer with 400 mM NaI instead of 140 mM NaCl) were added to achieve the extracellular I^-^ concentration of 100 mM. Further, forskolin / DMSO were added so that their concentration was not altered by the second fluid addition. After this, image acquisition continued for another 40 s.

### Image analysis

Images were analysed using MATLAB mathematical computing software (MathWorks) as described in (60). In brief, to estimate CFTR membrane proximity, a watershed transform-based segmentation was performed on binarized mCherry images after noise was removed. Cells were removed from analysis if they had (i) an area of <108 μm^2^ or >5400 μm^2^, (ii) a major axis length of >32.4 μm, (iii) if the area over the perimeter was <25 or >300 or (iv) if they were touching the image border. The membrane-proximal zone was defined as a 1.08 μm band within the border of each cell. After background correction, YFP and mCherry fluorescence intensity were normalized to the median YFP and mCherry fluorescence intensities of cells expressing WT CFTR on the same plate. Cells were removed from analysis if their average normalized fluorescence intensity fell below 0. For each cell, the CFTR membrane proximity (ρ) was defined as the average normalized YFP fluorescence intensity within the membrane-proximal zone over the average normalized mCherry fluorescence within the entire cell.

A separate image analysis protocol was used to assess CFTR activity. CFTR was activated with 10 μM forskolin and allowed to reach steady-state, after which 100 mM extracellular I^-^ was added. Images were corrected for background noise with help of a cell selection mask created using binarized mCherry images taken at the first and last timepoints. A mathematical model was then fitted to the cell YFP fluorescence time course (see below).

### mCherry fluorescence quality control

The mCherry fluorescence is related to the amount of bicistronic mRNA transcribed from the plasmid. The normalized mCherry fluorescence was quantified using images collected with the steady-state activity and membrane proximity protocols (see **Figure S10** and **Table S11**). In both datasets, mCherry fluorescence intensity of cells expressing F508del/Q1071F-CFTR was significantly lower compared to other variants. This difference persisted using a new DNA preparation of the plasmid. For this reason, the F508del/Q1071F mutant was not studied further and removed from the dataset.

### Mathematical model

The steady-state membrane potential (V_m_ in mV) and the CFTR-mediated whole-cell Cl^-^ conductance (G in nS) in the presence of 140 mM symmetrical [Cl^-^] can be estimated by fitting a mathematical model, with the experimental details defining initial conditions in the modelled system, to the fluorescence quenching time course (see (85)). A description of the model is provided in **Text S12**, and **Figure S13** shows the YFP(H148Q/I152L) quenching traces (solid red circles), together with the time course of several modelled variables, including the proportion of anion-free YFP(H148Q/I152L). Because, in our system, YFP(H148Q/I152L) fluorescence is completely quenched by anion binding (86), fitting the observed fluorescence time course to the predicted time course of the anion-free fluorophore allows parameter estimation. To account for variations in transfection efficiency, the mean mCherry fluorescence intensity within the cells was then used to normalize the G obtained by fitting.

At 28 °C, the estimated transient endogenous anion conductance (G_trans_) under control (DMSO) conditions decayed more slowly than previous estimates at 37 °C (85). We first ran the model with four free parameters (V_m_, G, G_trans_, and τ_trans_; see **Text S12**). However, at 28 °C there was more overlap between the transient and CFTR-mediated currents making it harder to reliably estimate the values of parameters describing the transient current. For this reason, we constrained G_trans_ and τ_trans_ to the average values obtained from the negative (DMSO) controls (9 nS and 11.4 s, respectively) and ran all the fits estimating only V_m_, and G.

### Molecular dynamics simulations

#### System setup

To build a starting structure for F508del-CFTR, we aligned one F508del-NBD1 from the human CFTR (hCFTR) F508del NBD1 dimer (PDB ID 2PZF; (64)) to the NBD1 of full-length ATP-bound, hCFTR (PDB ID 6MSM; (63)). This alignment was made based on the α- carbon atoms and gave a RMSD of 0.8 Å. A chimera model was then built by replacing the WT NBD1 with the F508del-NBD1 from PDB ID 2PZF. This system is referred to as F508del/R1070. ATP molecules and Mg^2+^ ions were included at the binding sites. We used this structure to prepare two additional simulation systems, namely F508del/R1070F and F508del/R1070W, by replacing the arginine at position 1070 with a phenylalanine and a tryptophan, respectively. This *in silico* mutagenesis was performed using PyMOL (PyMOL Molecular Graphics System, Version 2.5, Schrödinger, LLC.), and for each mutation, the rotamer with the least overlap with nearby residues was chosen.

System preparation for molecular dynamics (MD) simulations was performed using CHARMM-GUI (87– 89) as follows: Each F508del system was inserted in a POPC lipid bilayer using a simulation box of approximately 13, 13, and 18 nm in x, y, and z directions, respectively. This resulted in a total of 444 POPC lipids per system. Potassium and chloride ions were added at a concentration of 0.15 M. The simulations were performed with GROMACS 2019.3 (90, 91), using the CHARMM36 force field with the TIP3P water model, as implemented in GROMACS (92). Each system was minimized with position restraints on the backbone and side chain atoms using a force constant of 4000 and 2000 kJ·mol^-1^·nm^-2^, respectively, followed by multiple small equilibration steps to gradually decrease the force constant on the backbone atoms from 4000 to 0 kJ·mol^- 1^·nm^-2^, and from 2000 to 0 kJ·mol^-1^·nm^-2^for the side chain atoms, as recommended by the CHARMM-GUI protocol. The last stage of the equilibration was 20 ns long, with position restraints only on the backbone atoms using a force constant of 50 kJ·mol^-1^·nm^-2^. Production runs, with no position restraints applied, lasted 2 μs with a time step of 2 fs. The temperature was maintained at 303.15 K using the Nosé-Hoover algorithm (93, 94) and a relaxation time of 1 ps. Semi-isotropic pressure coupling was applied with the Parrinello-Rahman barostat (95) with a relaxation time of 12 ps, to maintain the pressure at 1 bar. The LINCS algorithm (96) was employed to constrain H-atoms bonds. Neighbour searching was made with the Verlet cutoff scheme, with van der Waals interactions switched to 0 between 1 and 1.2 nm. The long-range electrostatic interactions were treated using the Particle Mesh Ewald (PME) algorithm (97, 98).

#### Analyses

The cluster analysis was performed using the tool *gmx cluster* in GROMACS 2019.6. For this analysis, for each simulation system we extracted a trajectory containing only the atoms of the interface between ICL4 (residues 1050-1080 in hCFTR) and the F508del-NBD1 loop (residues 495-512 in hCFTR), with frames saved every 500 ps. The clustering was made with the Daura algorithm (99) using a cutoff of 0.1 nm. For the RMSD calculation, the alignment was performed on the backbone (N, αC, C) atoms of all the residues at the interface. The distribution of the χ1 dihedral angle (N-Cα-Cβ-Cγ) for selected aromatic residues at the interface was calculated using the *gmx angle* tool in GROMACS2019.6, and plotted using Matplotlib libraries (100). The probability density was calculated over 2 μs-long trajectories for the F508del systems, and over 1 μs-long trajectories for the previously published zCFTR systems (61). For clarity, although the ATP-bound protein is a Walker B glutamate E1372Q (corresponding to E1371Q in hCFTR) mutant, we refer to it here as “WT” to contrast it to the F508del mutants (note that the 6MSM hCFTR used to generate the F508del systems carries the same glutamate to glutamine mutation). The number of hydrogen bonds at the interface was estimated as a function of time based on a Hydrogen - Donor - Acceptor angle cutoff of 30 ° and on a Donor - Acceptor distance cutoff of 0.35 nm, as implemented in the *gmx hbond* tool in GROMACS2019.6. Plots were made using Matplotlib libraries (100). Figures with snapshots from simulations or experimental structures were generated with either VMD1.9.3 (101) or PyMOL 2.0.6 (The PyMOL Molecular Graphics System, Schrödinger, LLC).

### Data and statistical analysis

Statistical analysis was carried out in MATLAB (MathWorks) using the MATLAB Statistics Toolbox. Before statistical tests were performed, distributions were examined to assess whether they approximated normal distributions and whether there was homogeneity of variances. If the normality and homogeneity assumptions for parametric testing were not met, data were either transformed to meet the assumptions or analysed using non-parametric tests. When *post hoc* tests consisted of all possible pairwise comparisons between groups, the Tukey-Kramer procedure was applied (using the *multcompare* function in Matlab) to prevent inflation of the type I error rate. By contrast, when comparisons were planned, the Benjamini-Hochberg procedure with a false discovery rate of 10% was applied to control the family-wise error rate. To describe the data, the following commonly used statistical abbreviations were used: M (mean), Mdn (median), SD (standard deviation), SEM (standard error of the mean), N (number of measurements in the sample), and df (degrees of freedom).Statistical tests were performed two-sided unless otherwise specified. Data in the boxplots represent median, quartiles, range and outliers, and data in other graphs represent mean ± SEM. The significance level was pre-specified as α = 0.05. *, p < 0.05; **, p < 0.01; ***, p < 0.001; ****, p < 0.0001.

## Supporting information

description of mathematical model used for fitting quenching traces, tables detailing statistical analyses, additional figures

## Data availability

Most data are presented in the figures of the article. In addition, supporting information is available, which includes a description of the mathematical model used for fitting the quenching traces, tables detailing the statistical analyses performed, and additional figures illustrating the scan and molecular dynamics simulations

## Acknowledgments

We thank PT Harrison (University College Cork) and EA Miller (Cambridge University) for very interesting discussions. We also thank WD Andrews (Central Molecular Laboratory, University College London), and staff of the University College London Confocal Imaging Facility, for help with molecular biology and image acquisition, respectively.

## Author contributions

SP: Data Curation; Formal Analysis; Investigation; Software; Visualization; Writing – original draft; Writing – review & editing.

VC: Conceptualization; Data Curation; Formal Analysis; Investigation; Methodology; Software; Visualization; Writing – original draft; Writing – review & editing.

DNS: Funding Acquisition; Writing – review & editing.

DPT: Funding Acquisition; Resources; Writing – review & editing.

PV: Conceptualization; Funding Acquisition; Supervision; Writing – original draft; Writing – review & editing.

## Funding and additional information

SP, DNS and PV gratefully acknowledge funding by the Cystic Fibrosis Trust [SRC 005]. Work in DPT’s group is supported by the Natural Sciences and Engineering Research Council (Canada) and the Canadian Institutes of Health Research. Further support came from the Canada Research Chairs program. Calculations were carried out on Compute Canada facilities supported by the Canada Foundation for Innovation and partners.

## Conflict of interest

The authors declare that they have no conflicts of interest with the contents of this article.

### Abbreviations

The abbreviations used are

ABC: ATP-binding cassette
CFTR: cystic fibrosis transmembrane conductance regulator
G: whole-cell conductance
IRES: internal ribosome entry site
NBD: nucleotide-binding domain
P_o_: open probability
PDB: Protein Data Bank
TM1-12: transmembrane helix1 to 12
TMD: transmembrane domain
V_m_: membrane potential after steady-state activation of CFTR
YFP: yellow fluorescent protein.

## Notes

### Competing Interest Statement

The authors have declared no competing interest.

