## Supplementary material for "Can two wrongs make a right? F508del-CFTR ion channel rescue by second-site mutations in its transmembrane domains": description of mathematical model used for fitting quenching traces, tables detailing statistical analyses, additional figures

***Table S1*** **Descriptive statistics: assay readouts for WT CFTR and F508del-CFTR in the absence and presence of second-site mutations**

Colors highlight CFTR variants discussed in detail in text: F508del alone (yellow), or F508del in presence of F1068M (purple), of R1070W (blue), or of F1074M (red).

| | *Membrane proximity ($log_{10}\rho$)* | | | | *After addition of 10 µM forskolin* | | | | | | | *After addition of DMSO* | | | | | | |
|---|---|---|---|---|---|---|---|---|---|---|---|---|---|---|---|---|---|---|
| | | | | | | *G (nS)* | | | *$V_m$ (mV)* | | | | *G (nS)* | | | *$V_m$ (mV)* | | |
| | **N** | **M** | **Mdn** | **SD** | **N** | **M** | **Mdn** | **SD** | **M** | **Mdn** | **SD** | **N** | **M** | **Mdn** | **SD** | **M** | **Mdn** | **SD** |
| WT | 23 | -0.22 | -0.21 | 0.05 | 17 | 121.45 | 117.70 | 37.27 | -51.74 | -51.51 | 7.75 | 20 | 2.46 | 2.35 | 1.38 | -84.06 | -88.99 | 8.59 |
| F508del | 22 | -0.72 | -0.70 | 0.09 | 18 | 5.70 | 5.45 | 2.57 | -84.41 | -88.69 | 7.63 | 19 | 0.86 | 0.88 | 0.61 | -78.61 | -80.31 | *7.86* |
| F508del + | | | | | | | | | | | | | | | | | | |
| T1064F | 6 | -0.84 | -0.87 | 0.11 | 5 | 2.79 | 1.55 | 3.76 | -74.88 | -67.38 | 12.90 | 6 | 0.29 | 0.16 | 0.41 | -66.23 | -64.54 | *19.01* |
| T1064H | 6 | -0.74 | -0.72 | 0.10 | 5 | 7.05 | 1.48 | 8.47 | -74.03 | -74.88 | 14.81 | 5 | 0.23 | 0.20 | 0.29 | -58.65 | -53.93 | **15.79** |
| T1064M | 6 | -0.80 | -0.80 | 0.07 | 5 | 0.75 | 0.25 | 1.04 | -56.93 | -61.11 | 24.40 | 5 | 0.36 | 0.40 | 0.25 | -53.22 | -54.01 | 23.27 |
| T1064Q | 8 | -0.69 | -0.73 | 0.19 | 6 | 0.46 | 0.51 | 0.38 | -73.36 | -74.68 | 16.55 | 7 | 0.10 | 0.00 | 0.20 | -60.52 | -60.15 | 22.74 |
| T1064W | 6 | -0.80 | -0.78 | 0.09 | 5 | 0.18 | 0.14 | 0.17 | -51.68 | -48.27 | 17.53 | 5 | 0.53 | 0.43 | 0.56 | -68.47 | -57.93 | 19.64 |
| T1064Y | 6 | -0.78 | -0.76 | 0.13 | 5 | 3.04 | 0.57 | 5.63 | -61.93 | -64.97 | 21.71 | 5 | 0.08 | 0.00 | 0.17 | -58.03 | -56.13 | 15.31 |
| L1065F | 6 | -0.74 | -0.73 | 0.09 | 5 | 0.44 | 0.00 | 0.60 | -62.88 | -56.75 | 14.47 | 5 | 0.54 | 0.56 | 0.51 | -71.72 | -69.82 | 5.99 |
| L1065H | 6 | -0.77 | -0.74 | 0.09 | 5 | 0.77 | 0.43 | 0.73 | -71.14 | -77.83 | 14.19 | 5 | 0.60 | 0.60 | 0.43 | -63.95 | -63.20 | 10.64 |
| L1065M | 6 | -0.64 | -0.61 | 0.11 | 5 | 6.04 | 1.62 | 7.28 | -76.74 | -83.48 | 19.05 | 5 | 0.20 | 0.00 | 0.45 | -58.13 | -60.10 | 6.26 |
| L1065Q | 6 | -0.76 | -0.73 | 0.11 | 5 | 0.40 | 0.17 | 0.65 | -61.27 | -62.68 | 12.15 | 5 | 0.69 | 0.46 | 0.85 | -68.16 | -69.92 | 11.20 |
| L1065W | 6 | -0.81 | -0.83 | 0.10 | 5 | 0.60 | 0.36 | 0.46 | -73.38 | -80.84 | 18.25 | 5 | 0.91 | 1.04 | 0.37 | -84.15 | -83.02 | 5.53 |
| L1065Y | 11 | -0.77 | -0.75 | 0.09 | 10 | 0.76 | 0.36 | 0.78 | -69.75 | -68.40 | 17.06 | 10 | 0.71 | 0.45 | 0.64 | -81.71 | -79.77 | 7.60 |
| R1066F | 6 | -0.75 | -0.76 | 0.10 | 5 | 1.64 | 0.00 | 3.56 | -64.99 | -53.84 | 22.47 | 6 | 0.32 | 0.10 | 0.44 | -61.83 | -64.65 | 21.70 |
| R1066H | 6 | -0.76 | -0.76 | 0.04 | 5 | 2.63 | 0.32 | 5.42 | -60.28 | -62.77 | 20.22 | 5 | 1.66 | 0.97 | 2.59 | -67.75 | -76.12 | 25.28 |
| R1066M | 6 | -0.84 | -0.84 | 0.07 | 5 | 0.86 | 0.99 | 0.49 | -71.59 | -71.57 | 9.23 | 5 | 1.46 | 1.48 | 0.93 | -78.31 | -85.41 | 12.21 |
| R1066Q | 6 | -0.83 | -0.83 | 0.08 | 5 | 0.84 | 0.87 | 0.38 | -83.96 | -86.85 | 7.29 | 5 | 0.92 | 1.04 | 0.31 | -83.55 | -85.32 | 6.88 |
| R1066W | 11 | -0.85 | -0.89 | 0.17 | 9 | 0.85 | 1.00 | 0.69 | -71.64 | -76.23 | 15.47 | 9 | 0.85 | 0.66 | 0.87 | -74.92 | -80.91 | 17.46 |
| R1066Y | 6 | -0.80 | -0.78 | 0.06 | 5 | 0.11 | 0.05 | 0.15 | -49.29 | -42.53 | 20.36 | 5 | 0.25 | 0.00 | 0.53 | -53.18 | -51.20 | 10.24 |
| A1067F | 7 | -0.74 | -0.75 | 0.09 | 6 | 0.37 | 0.21 | 0.44 | -67.98 | -67.67 | 9.96 | 6 | 0.53 | 0.42 | 0.49 | -65.78 | -66.72 | 15.43 |
| A1067H | 7 | -0.75 | -0.73 | 0.09 | 6 | 3.15 | 0.96 | 5.10 | -67.84 | -66.91 | 16.00 | 6 | 0.46 | 0.27 | 0.64 | -65.98 | -71.13 | 12.44 |
| A1067M | 7 | -0.82 | -0.80 | 0.11 | 5 | 1.75 | 0.66 | 3.04 | -64.22 | -70.55 | 22.76 | 5 | 0.18 | 0.17 | 0.19 | -57.89 | -60.61 | 8.94 |
| A1067Q | 11 | -0.71 | -0.74 | 0.12 | 10 | 0.92 | 0.00 | 1.50 | -70.67 | -74.14 | 17.81 | 10 | 0.99 | 0.32 | 1.27 | -69.66 | -69.99 | 12.28 |
| A1067W | 6 | -0.80 | -0.82 | 0.07 | 5 | 1.06 | 1.18 | 0.34 | -71.69 | -75.10 | 8.24 | 5 | 0.45 | 0.50 | 0.34 | -75.63 | -74.61 | 9.19 |
| A1067Y | 6 | -0.81 | -0.82 | 0.10 | 5 | 0.63 | 0.50 | 0.69 | -70.10 | -71.79 | 15.95 | 5 | 0.35 | 0.29 | 0.16 | -68.46 | -63.01 | 9.32 |
| F1068H | 6 | -0.68 | -0.66 | 0.10 | 5 | 7.94 | 8.18 | 1.48 | -66.73 | -73.57 | 16.77 | 5 | 0.92 | 0.76 | 0.46 | -81.21 | -85.08 | 7.56 |
| F1068M | 6 | -0.70 | -0.65 | 0.12 | 5 | 18.25 | 20.39 | 6.28 | -58.02 | -55.26 | 9.07 | 5 | 0.48 | 0.48 | 0.35 | -72.71 | -71.90 | 12.94 |
| F1068Q | 6 | -0.74 | -0.70 | 0.13 | 5 | 11.27 | 8.89 | 6.38 | -68.50 | -69.00 | 15.44 | 5 | 0.79 | 0.71 | 0.44 | -75.69 | -73.92 | 10.51 |
| F1068W | 8 | -0.76 | -0.75 | 0.09 | 6 | 6.36 | 6.47 | 6.15 | -73.34 | -82.41 | 22.42 | 7 | 1.83 | 0.01 | 4.09 | -67.94 | -73.85 | 23.55 |
| F1068Y | 6 | -0.83 | -0.82 | 0.12 | 4 | 7.22 | 6.62 | 7.24 | -77.46 | -80.39 | 12.71 | 4 | 3.72 | 1.05 | 6.08 | -61.99 | -58.89 | 18.87 |
| G1069F | 10 | -0.88 | -0.89 | 0.09 | 10 | 8.63 | 0.33 | 22.69 | -66.90 | -68.68 | 16.98 | 10 | 0.63 | 0.49 | 0.66 | -72.16 | -68.15 | 12.32 |
| G1069H | 6 | -0.74 | -0.78 | 0.16 | 5 | 5.84 | 1.20 | 6.86 | -70.64 | -66.00 | 15.81 | 5 | 0.02 | 0.00 | 0.03 | -46.01 | -45.38 | 13.45 |
| G1069M | 6 | -0.81 | -0.85 | 0.11 | 5 | 2.09 | 2.21 | 1.09 | -87.21 | -87.47 | 3.09 | 5 | 1.26 | 1.30 | 0.72 | -82.34 | -89.75 | 10.32 |
| G1069Q | 11 | -0.73 | -0.74 | 0.15 | 10 | 0.62 | 0.64 | 0.45 | -60.81 | -61.86 | 14.84 | 10 | 0.54 | 0.33 | 0.76 | -61.45 | -64.88 | 18.14 |
| G1069W | 6 | -0.86 | -0.87 | 0.12 | 5 | 3.69 | 1.79 | 5.50 | -71.16 | -78.77 | 20.55 | 5 | 1.26 | 1.17 | 0.89 | -72.28 | -72.49 | 15.47 |
| G1069Y | 5 | -0.74 | -0.76 | 0.08 | 4 | 0.67 | 0.62 | 0.61 | -68.51 | -66.67 | 16.36 | 5 | 0.58 | 0.47 | 0.54 | -71.60 | -69.11 | 7.69 |
| R1070F | 6 | -0.74 | -0.76 | 0.10 | 5 | 8.73 | 8.71 | 1.02 | -68.03 | -70.21 | 12.76 | 5 | 1.68 | 1.57 | 0.50 | -75.56 | -73.29 | 9.30 |
| R1070H | 6 | -0.73 | -0.74 | 0.11 | 5 | 10.01 | 7.39 | 8.06 | -80.35 | -83.96 | 9.74 | 5 | 1.15 | 0.70 | 1.25 | -73.47 | -83.64 | 18.28 |
| R1070M | 6 | -0.56 | -0.56 | 0.14 | 5 | 7.83 | 7.93 | 1.75 | -56.11 | -61.95 | 20.66 | 5 | 0.42 | 0.41 | 0.33 | -71.55 | -73.34 | 14.22 |
| R1070Q | 11 | -0.78 | -0.75 | 0.10 | 10 | 11.37 | 10.36 | 6.19 | -73.72 | -74.61 | 8.41 | 11 | 0.78 | 0.46 | 0.73 | -69.09 | -68.11 | 7.80 |
| R1070W | 6 | -0.57 | -0.57 | 0.07 | 5 | 49.42 | 49.76 | 15.58 | -54.21 | -52.07 | 11.21 | 5 | 1.23 | 1.47 | 0.93 | -83.25 | -85.07 | 7.68 |
| R1070Y | 6 | -0.60 | -0.58 | 0.12 | 5 | 8.27 | 8.27 | 1.17 | -51.52 | -52.03 | 15.57 | 5 | 0.93 | 1.06 | 0.40 | -74.08 | -73.72 | 4.70 |
| Q1071H | 6 | -0.81 | -0.82 | 0.06 | 5 | 1.34 | 1.64 | 0.49 | -76.46 | -75.39 | 8.63 | 5 | 1.12 | 1.17 | 0.74 | -74.38 | -77.23 | 12.35 |
| Q1071M | 6 | -0.82 | -0.82 | 0.13 | 5 | 2.05 | 2.36 | 0.97 | -78.21 | -78.61 | 8.34 | 5 | 1.66 | 1.57 | 0.70 | -76.91 | -72.85 | 11.46 |
| Q1071W | 6 | -0.83 | -0.80 | 0.12 | 5 | 0.22 | 0.35 | 0.19 | -61.41 | -60.72 | 12.99 | 5 | 0.29 | 0.33 | 0.16 | -60.47 | -59.73 | 14.77 |
| Q1071Y | 6 | -0.80 | -0.79 | 0.06 | 5 | 0.40 | 0.35 | 0.35 | -51.01 | -40.23 | 15.69 | 5 | 0.17 | 0.00 | 0.25 | -51.20 | -51.02 | 11.74 |
| P1072F | 6 | -0.77 | -0.74 | 0.11 | 5 | 1.24 | 0.53 | 1.26 | -73.33 | -74.17 | 12.77 | 5 | 0.46 | 0.56 | 0.44 | -66.88 | -67.65 | 16.67 |
| P1072H | 7 | -0.79 | -0.87 | 0.18 | 6 | 7.17 | 7.12 | 7.06 | -78.34 | -81.35 | 11.17 | 6 | 1.97 | 0.63 | 3.62 | -75.93 | -76.27 | 9.82 |
| P1072M | 6 | -0.80 | -0.78 | 0.11 | 5 | 11.21 | 4.20 | 11.27 | -72.78 | -66.30 | 12.83 | 5 | 3.16 | 0.45 | 6.28 | -59.59 | -55.95 | 21.31 |
| P1072Q | 8 | -0.70 | -0.66 | 0.10 | 7 | 3.53 | 2.82 | 3.03 | -75.37 | -88.77 | 18.12 | 7 | 0.27 | 0.29 | 0.28 | -66.97 | -67.93 | 19.30 |
| P1072W | 6 | -0.87 | -0.88 | 0.11 | 5 | 8.36 | 8.15 | 4.49 | -81.58 | -85.79 | 12.60 | 5 | 1.12 | 1.31 | 0.49 | -76.44 | -75.51 | 7.89 |
| P1072Y | 6 | -0.77 | -0.77 | 0.16 | 5 | 9.95 | 7.60 | 6.46 | -85.49 | -86.91 | 4.40 | 5 | 0.81 | 0.57 | 1.00 | -73.83 | -77.71 | 14.76 |
| Y1073F | 6 | -0.82 | -0.85 | 0.17 | 5 | 8.13 | 1.81 | 9.13 | -70.26 | -65.00 | 15.02 | 5 | 9.54 | 11.3 | 9.47 | -77.35 | -85.11 | 13.22 |
| Y1073H | 6 | -0.70 | -0.68 | 0.13 | 5 | 3.82 | 2.51 | 2.88 | -74.97 | -79.50 | 12.30 | 5 | 0.95 | 0.89 | 0.71 | -79.13 | -80.40 | 11.05 |
| Y1073M | 6 | -0.74 | -0.71 | 0.10 | 5 | 3.70 | 1.72 | 5.06 | -70.47 | -78.09 | 21.15 | 5 | 0.49 | 0.01 | 0.78 | -72.22 | -69.10 | 13.27 |
| Y1073Q | 8 | -0.76 | -0.74 | 0.12 | 7 | 4.27 | 2.72 | 3.82 | -77.80 | -82.48 | 11.57 | 6 | 0.53 | 0.49 | 0.56 | -77.56 | -78.06 | 10.11 |
| Y1073W | 6 | -0.75 | -0.73 | 0.14 | 5 | 2.66 | 0.58 | 5.07 | -62.28 | -60.58 | 15.48 | 5 | 0.35 | 0.33 | 0.36 | -52.90 | -48.30 | 13.77 |
| F1074H | 6 | -0.80 | -0.77 | 0.16 | 5 | 0.46 | 0.19 | 0.60 | -52.69 | -45.66 | 17.99 | 5 | 0.30 | 0.23 | 0.26 | -55.67 | -53.42 | 12.81 |
| F1074M | 6 | -0.60 | -0.61 | 0.10 | 5 | 16.02 | 14.58 | 7.50 | -80.86 | -85.43 | 8.01 | 5 | 2.72 | 1.16 | 4.34 | -64.94 | -61.44 | 13.83 |
| F1074Q | 6 | -0.82 | -0.81 | 0.11 | 5 | 2.94 | 0.40 | 5.11 | -63.67 | -66.92 | 18.06 | 5 | 0.76 | 0.78 | 0.65 | -53.22 | -48.86 | 15.83 |
| F1074W | 6 | -0.84 | -0.86 | 0.14 | 4 | 4.21 | 2.64 | 4.66 | -71.09 | -78.87 | 21.94 | 5 | 1.31 | 1.30 | 0.97 | -70.15 | -68.47 | 13.99 |
| F1074Y | 6 | -0.78 | -0.75 | 0.11 | 5 | 1.03 | 0.61 | 0.95 | -53.73 | -54.00 | 8.67 | 5 | 0.37 | 0.44 | 0.34 | -61.38 | -61.04 | 9.72 |
| A141S | 6 | -0.80 | -0.74 | 0.19 | 5 | 16.17 | 14.29 | 9.35 | -74.40 | -73.51 | 13.77 | 4 | 0.39 | 0.43 | 0.36 | -69.52 | -70.20 | 17.71 |
| R1097T | 6 | -0.71 | -0.69 | 0.09 | 5 | 15.30 | 13.46 | 9.25 | -68.79 | -70.24 | 16.98 | 5 | 0.32 | 0.02 | 0.49 | -71.56 | -70.75 | 11.89 |

***Table S2* One-tailed Wilcoxon Rank Sum tests (DMSO vs. forskolin)**

For every mutant the test assessed whether conductance (G) was significantly increased and/or whether the membrane potential ($V_m$) was significantly depolarized after addition of 10 µM forskolin compared to the DMSO control condition. The colors highlight the CFTR variants discussed. W indicates the Wilcoxon rank-sum test statistic, z the z-score, and P the p-value.

| | *G* | | | | *$V_m$* | | | |
|---|---|---|---|---|---|---|---|---|
| | **W** | **z** | **P** | | **W** | **z** | **P** | |
| WT | 210 | -5.17 | 1.20E-07 | **** | 211 | -5.14 | 1.41E-07 | **** |
| F508del | 206 | -4.69 | 1.33E-06 | **** | 426 | 1.99 | 0.977 | |
| F508del + T1064F | 25 | -1.92 | 0.028 | * | 41 | 1.00 | 0.842 | |
| F508del + T1064H | 16 | -2.30 | 0.011 | * | 34 | 1.46 | 0.928 | |
| F508del + T1064M | 27 | 0.00 | 0.500 | | 29 | 0.42 | 0.662 | |
| F508del + T1064Q | 37 | -1.64 | 0.050 | | 56 | 1.07 | 0.858 | |
| F508del + T1064W | 31 | 0.63 | 0.798 | | 21 | -1.25 | 0.105 | |
| F508del + T1064Y | 17 | -2.09 | 0.018 | * | 29 | 0.42 | 0.662 | |
| F508del + L1065F | 32 | 0.84 | 0.852 | | 23 | -0.84 | 0.202 | |
| F508del + L1065H | 26 | -0.21 | 0.417 | | 33 | 1.25 | 0.895 | |
| F508del + L1065M | 17 | -2.09 | 0.018 | * | 35 | 1.67 | 0.953 | |
| F508del + L1065Q | 33 | 1.04 | 0.895 | | 25 | -0.42 | 0.338 | |
| F508del + L1065W | 33 | 1.04 | 0.895 | | 23 | -0.84 | 0.202 | |
| F508del + L1065Y | 109 | 0.26 | 0.633 | | 84 | -1.55 | 0.061 | |
| F508del + R1066F | 37 | 0.09 | 0.608 | | 39 | 0.64 | 0.739 | |
| F508del + R1066H | 29 | 0.21 | 0.662 | | 24 | -0.63 | 0.265 | |
| F508del + R1066M | 35 | 1.46 | 0.953 | | 21 | -1.25 | 0.105 | |
| F508del + R1066Q | 30 | 0.42 | 0.735 | | 29 | 0.42 | 0.662 | |
| F508del + R1066W | 82 | -0.26 | 0.396 | | 74 | -0.97 | 0.166 | |
| F508del + R1066Y | 26 | -0.21 | 0.417 | | 24 | -0.63 | 0.265 | |
| F508del + A1067F | 44 | 0.72 | 0.811 | | 38 | -0.08 | 0.468 | |
| F508del + A1067H | 30 | -1.36 | 0.087 | | 39 | 0.08 | 0.532 | |
| F508del + A1067M | 21 | -1.25 | 0.105 | | 30 | 0.63 | 0.735 | |
| F508del + A1067Q | 120 | 1.10 | 0.879 | | 108 | 0.26 | 0.604 | |
| F508del + A1067W | 17 | -2.09 | 0.018 | * | 27 | 0.00 | 0.500 | |
| F508del + A1067Y | 26 | -0.21 | 0.417 | | 28 | 0.21 | 0.583 | |
| F508del + F1068H | 15 | -2.51 | 0.006 | ** | 20 | -1.46 | 0.072 | |
| F508del + F1068M | 15 | -2.51 | 0.006 | ** | 18 | -1.88 | 0.030 | * |
| F508del + F1068Q | 15 | -2.51 | 0.006 | ** | 24 | -0.63 | 0.265 | |
| F508del + F1068W | 39 | -1.36 | 0.087 | | 50 | 0.21 | 0.585 | |
| F508del + F1068Y | 15 | -0.72 | 0.235 | | 23 | 1.59 | 0.944 | |
| F508del + G1069F | 108 | 0.19 | 0.604 | | 97 | -0.57 | 0.285 | |
| F508del + G1069H | 15 | -2.51 | 0.006 | ** | 37 | 2.09 | 0.982 | |
| F508del + G1069M | 21 | -1.25 | 0.105 | | 27 | 0.00 | 0.500 | |
| F508del + G1069Q | 92 | -0.94 | 0.172 | | 103 | -0.11 | 0.455 | |
| F508del + G1069W | 26 | -0.21 | 0.417 | | 27 | 0.00 | 0.500 | |
| F508del + G1069Y | 25 | 0.00 | 0.549 | | 23 | -0.37 | 0.357 | |
| F508del + R1070F | 15 | -2.51 | 0.006 | ** | 24 | -0.63 | 0.265 | |
| F508del + R1070H | 15 | -2.51 | 0.006 | ** | 32 | 1.04 | 0.852 | |
| F508del + R1070M | 15 | -2.51 | 0.006 | ** | 23 | -0.84 | 0.202 | |
| F508del + R1070Q | 66 | -3.84 | 6.21E-05 | **** | 138 | 1.23 | 0.891 | |
| F508del + R1070W | 15 | -2.51 | 0.006 | ** | 15 | -2.51 | 0.006 | ** |
| F508del + R1070Y | 15 | -2.51 | 0.006 | ** | 16 | -2.30 | 0.011 | * |
| F508del + Q1071H | 24 | -0.63 | 0.265 | | 27 | 0.00 | 0.500 | |
| F508del + Q1071M | 25 | -0.42 | 0.338 | | 28 | 0.21 | 0.583 | |
| F508del + Q1071W | 27 | 0.00 | 0.500 | | 28 | 0.21 | 0.583 | |
| F508del + Q1071Y | 20 | -1.46 | 0.072 | | 27 | 0.00 | 0.500 | |
| F508del + P1072F | 24 | -0.63 | 0.265 | | 30 | 0.63 | 0.735 | |
| F508del + P1072H | 33 | -0.88 | 0.189 | | 42 | 0.56 | 0.712 | |
| F508del + P1072M | 18 | -1.88 | 0.030 | * | 32 | 1.04 | 0.852 | |
| F508del + P1072Q | 38 | -1.79 | 0.037 | * | 61 | 1.15 | 0.875 | |
| F508del + P1072W | 15 | -2.51 | 0.006 | ** | 32 | 1.04 | 0.852 | |
| F508del + P1072Y | 15 | -2.51 | 0.006 | ** | 34 | 1.46 | 0.928 | |
| F508del + Y1073F | 26 | -0.21 | 0.417 | | 22 | -1.04 | 0.148 | |
| F508del + Y1073H | 18 | -1.88 | 0.030 | * | 25 | -0.42 | 0.338 | |
| F508del + Y1073M | 21 | -1.25 | 0.105 | | 26 | -0.21 | 0.417 | |
| F508del + Y1073Q | 27 | -2.07 | 0.019 | * | 43 | 0.21 | 0.585 | |
| F508del + Y1073W | 24 | -0.63 | 0.265 | | 31 | 0.84 | 0.798 | |
| F508del + F1074H | 26 | -0.21 | 0.417 | | 26 | -0.21 | 0.417 | |
| F508del + F1074M | 16 | -2.30 | 0.011 | * | 35 | 1.67 | 0.953 | |
| F508del + F1074Q | 26 | -0.21 | 0.417 | | 32 | 1.04 | 0.852 | |
| F508del + F1074W | 20 | -1.10 | 0.135 | | 26 | 0.37 | 0.643 | |
| F508del + F1074Y | 21 | -1.25 | 0.105 | | 22 | -1.04 | 0.148 | |
| F508del + A141S | 10 | -2.33 | 0.010 | ** | 21 | 0.37 | 0.643 | |
| F508del + R1097T | 15 | -2.51 | 0.006 | ** | 27 | 0.00 | 0.500 | |

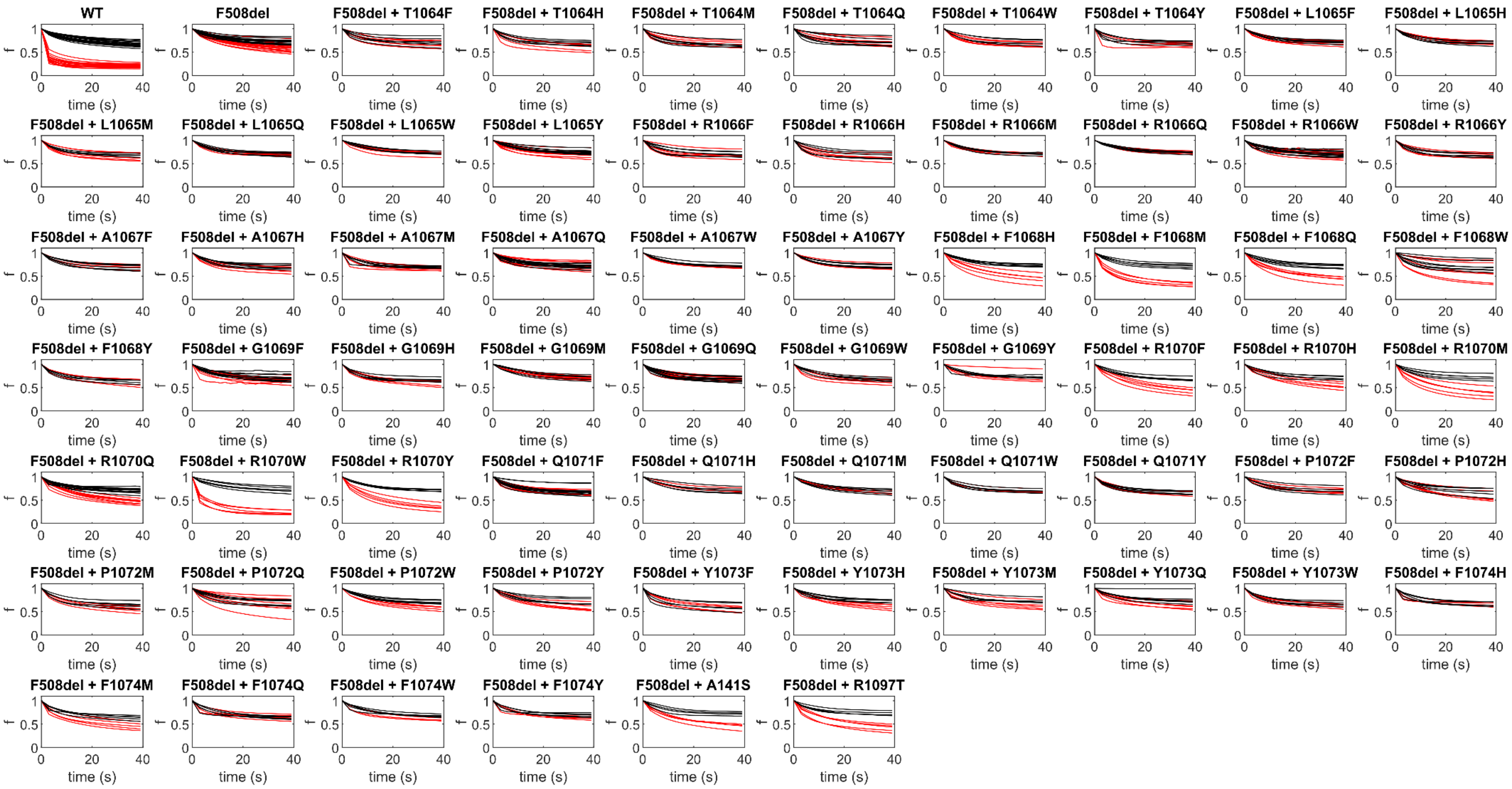


***Figure S3*** **Fluorescence quenching traces**

Fluorescence quenching timelines measured in HEK-293 cells expressing WT CFTR or F508del-CFTR in the absence or presence of second-site mutations. 230 s after addition of 10 µM forskolin (red) or DMSO (black), $I^-$ was added at time point 0. The fluorescence (f) was normalized to the timepoint before $I^-$ addition.

***Table S4*** **Two-tailed Wilcoxon Rank Sum tests comparing ion channel function of cells expressing F508del in the absence and presence of second-site mutations**

Comparison of median G and $V_m$ after addition of 10 µM forskolin to HEK-293 cells expressing F508del-CFTR and F508del with second-site mutations. The table shows the Wilcoxon rank-sum test statistic (W), and the z-score (z). The Benjamini-Hochberg procedure with a false discovery rate of 10% was applied to control the family wise error rate. P-values (P) below the critical (Q) value were considered significant (above the dotted lines). Note that in many cases the second-site mutation causes a reduction in G. The colors highlight F508del together with F1068M (purple), R1070W (blue), and F1074M (red).

| | *G* | | | | | *$V_m$* | | | |
|---|---|---|---|---|---|---|---|---|---|
| | **W** | **z** | **P** | **Q** | | **W** | **z** | **P** | **Q** |
| F508del + L1065Y | 343 | 3.91 | 9.32E-05 | 0.002 | F508del + G1069Q | 179 | -3.91 | 9.32E-05 | 0.002 |
| F508del + G1069Q | 343 | 3.91 | 9.32E-05 | 0.003 | F508del + R1070W | 171 | -3.32 | 0.001 | 0.003 |
| F508del + R1066W | 327 | 3.83 | 1.27E-04 | 0.005 | F508del + F1074Y | 171 | -3.32 | 0.001 | 0.005 |
| F508del + A1067Q | 341 | 3.81 | 1.38E-04 | 0.006 | F508del + G1069F | 193 | -3.24 | 0.001 | 0.006 |
| F508del + A1067F | 277 | 3.43 | 0.001 | 0.008 | F508del + F1068M | 174 | -3.09 | 0.002 | 0.008 |
| F508del + R1070W | 171 | -3.32 | 0.001 | 0.010 | F508del + R1070Y | 174 | -3.09 | 0.002 | 0.010 |
| F508del + T1064Q | 275 | 3.30 | 0.001 | 0.011 | F508del + Q1071Y | 174 | -3.09 | 0.002 | 0.011 |
| F508del + T1064W | 260 | 3.24 | 0.001 | 0.013 | F508del + T1064W | 175 | -3.02 | 0.003 | 0.013 |
| F508del + R1066Y | 260 | 3.24 | 0.001 | 0.014 | F508del + R1066Y | 175 | -3.02 | 0.003 | 0.014 |
| F508del + L1065F | 259 | 3.17 | 0.002 | 0.016 | F508del + R1070M | 176 | -2.94 | 0.003 | 0.016 |
| F508del + L1065Q | 259 | 3.17 | 0.002 | 0.017 | F508del + Q1071W | 176 | -2.94 | 0.003 | 0.017 |
| F508del + F1074H | 259 | 3.17 | 0.002 | 0.019 | F508del + F1074H | 176 | -2.94 | 0.003 | 0.019 |
| F508del + T1064M | 258 | 3.09 | 0.002 | 0.021 | F508del + R1070Q | 200 | -2.90 | 0.004 | 0.021 |
| F508del + L1065W | 258 | 3.09 | 0.002 | 0.022 | F508del + L1065Q | 177 | -2.87 | 0.004 | 0.022 |
| F508del + A1067Y | 258 | 3.09 | 0.002 | 0.024 | F508del + Y1073W | 177 | -2.87 | 0.004 | 0.024 |
| F508del + Q1071W | 258 | 3.09 | 0.002 | 0.025 | F508del + A1067F | 182 | -2.83 | 0.005 | 0.025 |
| F508del + Q1071Y | 258 | 3.09 | 0.002 | 0.027 | F508del + R1066H | 178 | -2.80 | 0.005 | 0.027 |
| F508del + F1074M | 174 | -3.09 | 0.002 | 0.029 | F508del + A1067H | 183 | -2.77 | 0.006 | 0.029 |
| F508del + L1065H | 257 | 3.02 | 0.003 | 0.030 | F508del + L1065F | 179 | -2.72 | 0.007 | 0.030 |
| F508del + R1066M | 257 | 3.02 | 0.003 | 0.032 | F508del + R1066W | 199 | -2.70 | 0.007 | 0.032 |
| F508del + R1066Q | 256 | 2.94 | 0.003 | 0.033 | F508del + F1074Q | 180 | -2.65 | 0.008 | 0.033 |
| F508del + A1067W | 256 | 2.94 | 0.003 | 0.035 | F508del + T1064Y | 181 | -2.57 | 0.010 | 0.035 |
| F508del + F1068M | 176 | -2.94 | 0.003 | 0.037 | F508del + R1070F | 181 | -2.57 | 0.010 | 0.037 |
| F508del + Q1071H | 256 | 2.94 | 0.003 | 0.038 | F508del + F1068Q | 182 | -2.50 | 0.013 | 0.038 |
| F508del + F1074Y | 256 | 2.94 | 0.003 | 0.040 | F508del + R1066M | 183 | -2.42 | 0.015 | 0.040 |
| F508del + P1072F | 255 | 2.87 | 0.004 | 0.041 | F508del + A1067W | 183 | -2.42 | 0.015 | 0.041 |
| F508del + R1097T | 178 | -2.80 | 0.005 | 0.043 | F508del + F1068H | 183 | -2.42 | 0.015 | 0.043 |
| F508del + G1069Y | 240 | 2.77 | 0.006 | 0.044 | F508del + G1069H | 183 | -2.42 | 0.015 | 0.044 |
| F508del + G1069M | 253 | 2.72 | 0.007 | 0.046 | F508del + P1072M | 183 | -2.42 | 0.015 | 0.046 |
| F508del + Q1071M | 253 | 2.72 | 0.007 | 0.048 | F508del + Y1073F | 183 | -2.42 | 0.015 | 0.048 |
| F508del + A141S | 179 | -2.72 | 0.007 | 0.049 | F508del + R1097T | 183 | -2.42 | 0.015 | 0.049 |
| F508del + R1070Q | 208 | -2.52 | 0.012 | 0.051 | F508del + A1067M | 184 | -2.35 | 0.019 | 0.051 |
| F508del + F1068Q | 183 | -2.42 | 0.015 | 0.052 | F508del + L1065H | 185 | -2.27 | 0.023 | 0.052 |
| F508del + G1069F | 311 | 2.37 | 0.018 | 0.054 | F508del + T1064M | 188 | -2.05 | 0.040 | 0.054 |
| F508del + R1066F | 248 | 2.35 | 0.019 | 0.056 | F508del + Y1073Q | 200 | -2.03 | 0.043 | 0.056 |
| F508del + R1070F | 185 | -2.27 | 0.023 | 0.057 | F508del + G1069W | 189 | -1.98 | 0.048 | 0.057 |
| F508del + A1067M | 246 | 2.20 | 0.028 | 0.059 | F508del + Y1073H | 189 | -1.98 | 0.048 | 0.059 |
| F508del + A1067H | 256 | 2.03 | 0.042 | 0.060 | F508del + A141S | 189 | -1.98 | 0.048 | 0.060 |
| F508del + R1066H | 241 | 1.83 | 0.068 | 0.062 | F508del + T1064H | 190 | -1.90 | 0.057 | 0.062 |
| F508del + R1070Y | 191 | -1.83 | 0.068 | 0.063 | F508del + F1074W | 185 | -1.83 | 0.067 | 0.063 |
| F508del + T1064F | 240 | 1.75 | 0.080 | 0.065 | F508del + F1068Y | 186 | -1.75 | 0.081 | 0.065 |
| F508del + T1064Y | 240 | 1.75 | 0.080 | 0.067 | F508del + L1065M | 193 | -1.68 | 0.094 | 0.067 |
| F508del + Y1073W | 240 | 1.75 | 0.080 | 0.068 | F508del + T1064F | 194 | -1.60 | 0.109 | 0.068 |
| F508del + F1074Q | 240 | 1.75 | 0.080 | 0.070 | F508del + R1066F | 195 | -1.53 | 0.127 | 0.070 |
| F508del + F1068H | 193 | -1.68 | 0.094 | 0.071 | F508del + P1072F | 195 | -1.53 | 0.127 | 0.071 |
| F508del + G1069W | 239 | 1.68 | 0.094 | 0.073 | F508del + F1074M | 195 | -1.53 | 0.127 | 0.073 |
| F508del + R1070M | 193 | -1.68 | 0.094 | 0.075 | F508del + L1065Y | 229 | -1.51 | 0.131 | 0.075 |
| F508del + P1072Q | 259 | 1.48 | 0.138 | 0.076 | F508del + A1067Y | 196 | -1.45 | 0.146 | 0.076 |
| F508del + P1072Y | 196 | -1.45 | 0.146 | 0.078 | F508del + G1069Y | 190 | -1.40 | 0.160 | 0.078 |
| F508del + Y1073H | 236 | 1.45 | 0.146 | 0.079 | F508del + L1065W | 198 | -1.30 | 0.192 | 0.079 |
| F508del + Y1073M | 236 | 1.45 | 0.146 | 0.081 | F508del + Y1073M | 198 | -1.30 | 0.192 | 0.081 |
| F508del + P1072W | 200 | -1.16 | 0.248 | 0.083 | F508del + A1067Q | 234 | -1.27 | 0.204 | 0.083 |
| F508del + R1070H | 201 | -1.08 | 0.280 | 0.084 | F508del + P1072H | 206 | -1.23 | 0.217 | 0.084 |
| F508del + F1074W | 220 | 1.06 | 0.287 | 0.086 | F508del + Q1071M | 199 | -1.23 | 0.219 | 0.086 |
| F508del + Y1073Q | 251 | 1.00 | 0.318 | 0.087 | F508del + Q1071H | 200 | -1.16 | 0.248 | 0.087 |
| F508del + T1064H | 222 | 0.41 | 0.682 | 0.089 | F508del + R1070H | 201 | -1.08 | 0.280 | 0.089 |
| F508del + L1065M | 222 | 0.41 | 0.682 | 0.090 | F508del + P1072W | 202 | -1.01 | 0.314 | 0.090 |
| F508del + G1069H | 222 | 0.41 | 0.682 | 0.092 | F508del + P1072Y | 204 | -0.86 | 0.391 | 0.092 |
| F508del + Y1073F | 222 | 0.41 | 0.682 | 0.094 | F508del + T1064Q | 212 | -0.83 | 0.405 | 0.094 |
| F508del + P1072H | 220 | -0.30 | 0.764 | 0.095 | F508del + F1068W | 216 | -0.57 | 0.571 | 0.095 |
| F508del + F1068Y | 206 | -0.04 | 0.966 | 0.097 | F508del + G1069M | 221 | 0.34 | 0.737 | 0.097 |
| F508del + P1072M | 215 | -0.04 | 0.970 | 0.098 | F508del + R1066Q | 218 | 0.11 | 0.911 | 0.098 |
| F508del + F1068W | 224 | -0.03 | 0.973 | 0.100 | F508del + P1072Q | 232 | -0.09 | 0.928 | 0.100 |

***Table S5 P*aired t-tests comparing $log_{10}\rho$ of F508del with F508del and second-site mutations**

Statistical tests were performed on the difference between $log_{10}\rho$ obtained for HEK-293 cells expressing F508del-CFTR in the absence and presence of second-site mutations, from the same plate (mean, M, and standard deviation, SD, of this difference are shown in leftmost columns). The Benjamini-Hochberg procedure with a false discovery rate of 10% was applied to control for the family wise error rate. P-values (P) below the critical value (Q) are considered significant (above dotted line). Again, many second-site mutations worsen the F508del defect. T is the T-value and the colors highlight variants most discussed in text.

| | *Difference* | | *Test statistics* | | | |
|---|---|---|---|---|---|---|
| | **M** | **SD** | **T** | **df** | **P** | **Q** |
| F508del + Q1071H | -0.13 | 0.05 | 6.44 | 5 | 0.001 | 0.002 |
| F508del + R1066M | -0.15 | 0.06 | 5.81 | 5 | 0.002 | 0.003 |
| F508del + F1068Y | -0.14 | 0.06 | 5.79 | 5 | 0.002 | 0.005 |
| F508del + A1067F | -0.06 | 0.03 | 4.86 | 6 | 0.003 | 0.006 |
| F508del + R1066Y | -0.11 | 0.05 | 5.26 | 5 | 0.003 | 0.008 |
| F508del + G1069F | -0.15 | 0.12 | 3.94 | 9 | 0.003 | 0.010 |
| F508del + R1066Q | -0.14 | 0.07 | 5.05 | 5 | 0.004 | 0.011 |
| F508del + A1067Y | -0.12 | 0.06 | 5.04 | 5 | 0.004 | 0.013 |
| F508del + P1072W | -0.20 | 0.10 | 4.84 | 5 | 0.005 | 0.014 |
| F508del + A1067M | -0.09 | 0.05 | 4.34 | 6 | 0.005 | 0.016 |
| F508del + T1064W | -0.11 | 0.06 | 4.56 | 5 | 0.006 | 0.017 |
| F508del + R1066H | -0.10 | 0.06 | 4.37 | 5 | 0.007 | 0.019 |
| F508del + R1070W | 0.15 | 0.09 | -4.26 | 5 | 0.008 | 0.021 |
| F508del + Q1071Y | -0.09 | 0.05 | 4.21 | 5 | 0.008 | 0.022 |
| F508del + T1064M | -0.15 | 0.09 | 3.97 | 5 | 0.011 | 0.024 |
| F508del + F1074Q | -0.09 | 0.05 | 4.11 | 4 | 0.015 | 0.025 |
| F508del + L1065W | -0.12 | 0.08 | 3.55 | 5 | 0.016 | 0.027 |
| F508del + F1074M | 0.15 | 0.08 | -3.98 | 4 | 0.016 | 0.029 |
| F508del + L1065Y | -0.06 | 0.07 | 2.87 | 10 | 0.017 | 0.030 |
| F508del + A1067H | -0.07 | 0.06 | 3.22 | 6 | 0.018 | 0.032 |
| F508del + L1065F | -0.07 | 0.05 | 3.39 | 5 | 0.019 | 0.033 |
| F508del + A1067W | -0.11 | 0.08 | 3.25 | 5 | 0.023 | 0.035 |
| F508del + P1072M | -0.13 | 0.10 | 3.20 | 5 | 0.024 | 0.037 |
| F508del + R1070M | 0.16 | 0.13 | -3.09 | 5 | 0.027 | 0.038 |
| F508del + P1072F | -0.10 | 0.08 | 2.95 | 5 | 0.032 | 0.040 |
| F508del + Q1071W | -0.11 | 0.09 | 2.92 | 5 | 0.033 | 0.041 |
| F508del + G1069W | -0.14 | 0.12 | 2.91 | 5 | 0.033 | 0.043 |
| F508del + Q1071M | -0.14 | 0.13 | 2.79 | 5 | 0.038 | 0.044 |
| F508del + R1070Y | 0.12 | 0.12 | -2.58 | 5 | 0.049 | 0.046 |
| F508del + F1068W | -0.07 | 0.08 | 2.36 | 7 | 0.051 | 0.048 |
| F508del + L1065Q | -0.07 | 0.07 | 2.52 | 5 | 0.053 | 0.049 |
| F508del + T1064F | -0.09 | 0.09 | 2.49 | 5 | 0.055 | 0.051 |
| F508del + L1065M | 0.05 | 0.05 | -2.47 | 5 | 0.057 | 0.052 |
| F508del + Y1073F | -0.15 | 0.16 | 2.32 | 5 | 0.068 | 0.054 |
| F508del + G1069M | -0.09 | 0.10 | 2.30 | 5 | 0.070 | 0.056 |
| F508del + T1064H | -0.05 | 0.06 | 2.28 | 5 | 0.072 | 0.057 |
| F508del + L1065H | -0.08 | 0.08 | 2.28 | 5 | 0.072 | 0.059 |
| F508del + R1066W | -0.11 | 0.19 | 1.86 | 10 | 0.093 | 0.060 |
| F508del + T1064Y | -0.05 | 0.07 | 1.78 | 5 | 0.135 | 0.062 |
| F508del + R1070Q | -0.06 | 0.12 | 1.57 | 10 | 0.147 | 0.063 |
| F508del + P1072H | -0.09 | 0.15 | 1.66 | 6 | 0.147 | 0.065 |
| F508del + R1066F | -0.09 | 0.13 | 1.71 | 5 | 0.148 | 0.067 |
| F508del + A141S | -0.11 | 0.16 | 1.66 | 5 | 0.157 | 0.068 |
| F508del + P1072Y | -0.10 | 0.16 | 1.56 | 5 | 0.179 | 0.070 |
| F508del + F1074W | -0.10 | 0.15 | 1.60 | 4 | 0.185 | 0.071 |
| F508del + Y1073W | -0.08 | 0.13 | 1.47 | 5 | 0.201 | 0.073 |
| F508del + F1074Y | -0.05 | 0.07 | 1.40 | 4 | 0.233 | 0.075 |
| F508del + F1068H | 0.04 | 0.08 | -1.26 | 5 | 0.264 | 0.076 |
| F508del + Y1073Q | -0.04 | 0.08 | 1.19 | 6 | 0.278 | 0.078 |
| F508del + F1074H | -0.05 | 0.10 | 1.13 | 4 | 0.322 | 0.079 |
| F508del + R1097T | -0.02 | 0.05 | 1.10 | 5 | 0.322 | 0.081 |
| F508del + F1068Q | -0.02 | 0.09 | 0.69 | 5 | 0.520 | 0.083 |
| F508del + F1068M | 0.02 | 0.08 | -0.66 | 5 | 0.538 | 0.084 |
| F508del + A1067Q | 0.03 | 0.15 | -0.60 | 10 | 0.563 | 0.086 |
| F508del + G1069H | -0.03 | 0.13 | 0.48 | 5 | 0.649 | 0.087 |
| F508del + G1069Y | -0.02 | 0.11 | 0.48 | 4 | 0.655 | 0.089 |
| F508del + R1070F | -0.02 | 0.10 | 0.45 | 5 | 0.670 | 0.090 |
| F508del + Y1073H | 0.01 | 0.08 | -0.41 | 4 | 0.700 | 0.092 |
| F508del + P1072Q | -0.01 | 0.08 | 0.38 | 7 | 0.713 | 0.094 |
| F508del + Y1073M | -0.01 | 0.09 | 0.21 | 4 | 0.843 | 0.095 |
| F508del + R1070H | -0.01 | 0.11 | 0.20 | 5 | 0.847 | 0.097 |
| F508del + T1064Q | 0.01 | 0.20 | -0.08 | 7 | 0.938 | 0.098 |
| F508del + G1069Q | 0.00 | 0.15 | 0.06 | 10 | 0.957 | 0.100 |

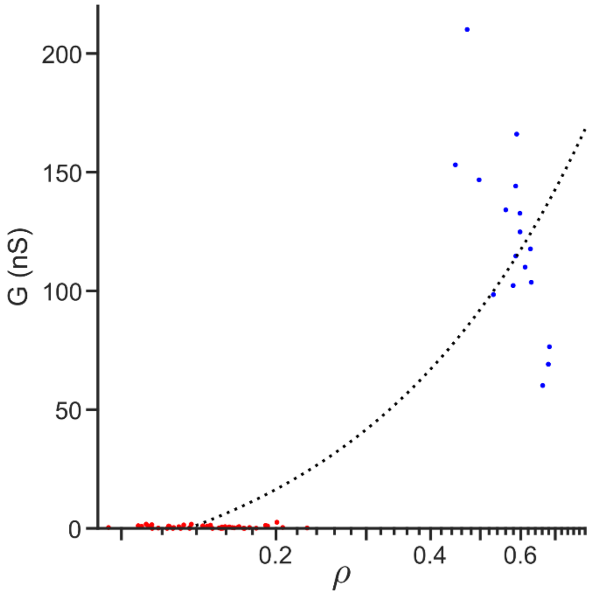


***Figure S6*** **G-ρ relationship**

To estimate the ρ value at which there are virtually no channels at the plasma membrane, the most impaired mutants – with an average G < 1 nS and an average $\log_{10}\rho < -0.8$ – were selected from the screen (T1064M, R1066M, R1066Q, R1066W, R1066Y, R1067Y, Q1071W, Q1071Y; red filled circles). The average ρ value of these mutants was 0.15. A restrained linear regression was performed on the G-ρ measurements of WT CFTR after basal activation with 10 μM forskolin (blue filled circles), forcing the regression through the x-axis intercept at ρ = 0.15. G was plotted as a function of membrane proximity (ρ, obtained by back transformation of mean $\log_{10}\rho$).

***Table S7* ICL4/NBD1 and ICL2/NBD2 interface**

Residues relevant to the analyses of the MD simulations, forming the ICL4/NBD1 interface and the ICL2/NBD2 interface in human CFTR (hCFTR) and zebrafish CFTR (zCFTR). For the ICL4/NBD1 interface the complete alignment is shown below.

| | **hCFTR** | **zCFTR** |
|---|---|---|
| **ICL4** | F1068 | F1076 |
| | R1070 | R1078 |
| | F1074 | F1082 |
| **NBD1 loop** | E504 | D503 |
| | I507 | L506 |
| | F508 | F507 |
| | G509 | G508 |
| **ICL2** | Y275 | Y276 |
| | W277 | W278 |
| | M281 | M282 |
| **NBD2 loop** | P1306 | P1307 |

**NBD1 loop**

| | | | |
|---|---|---|---|
| human | 495 | SWIMPGTIKENIIFGVSY | 512 |
| zebrafish | 494 | AWIMPGTIRDNILFGLTY | 511 |

**ICL4**

| | | | |
|---|---|---|---|
| human | 1050 | PIFTHLVTSLKGLWTLRAFGRQPYFETLFHK | 1080 |
| zebrafish | 1058 | PIFSHLIMSLKGLWTIRAFERQAYFEALFHK | 1088 |

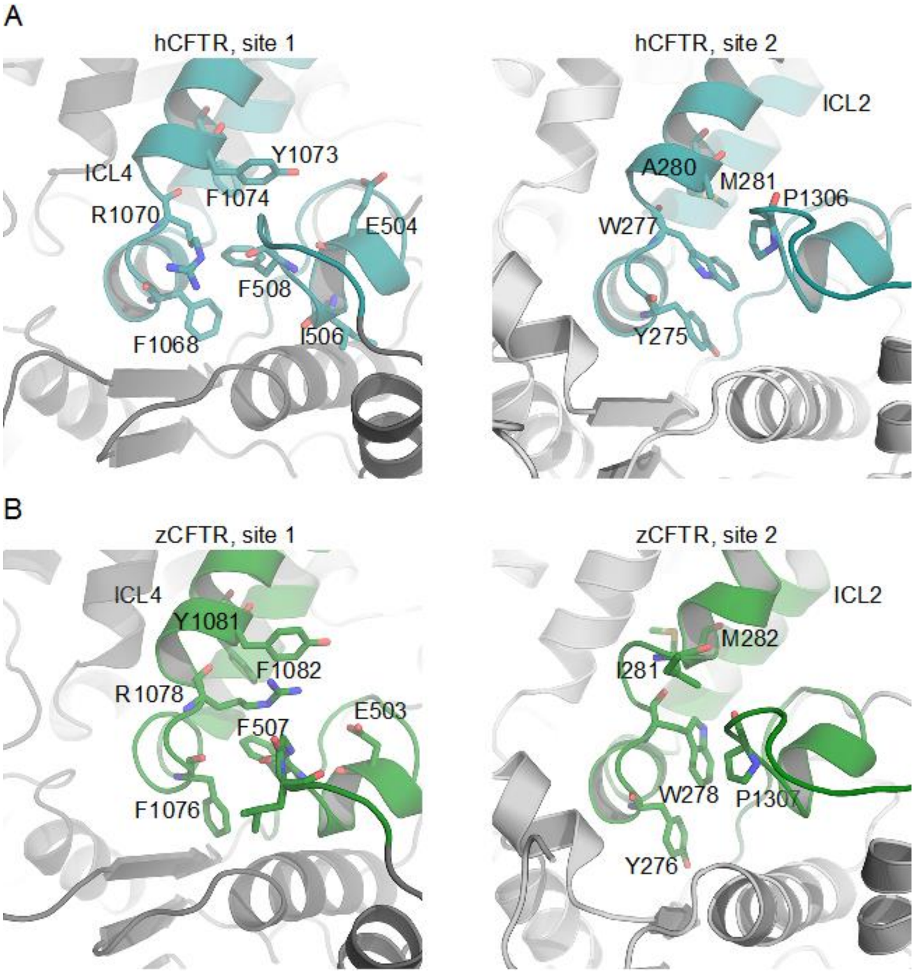


***Figure S8*** **The interface between ICL4 and NBD1 and between ICL2 and NBD2 in human CFTR (hCFTR) and zebrafish CFTR (zCFTR)**

**A**) Left, residues 1050-1080 of ICL4 and residues 495-511 of NBD1 and right, residues 256-286 of ICL2 and residues 1294-1311 of NBD2 of hCFTR (PDB ID 6MSM, (9)) are shown in cyan cartoons. Selected residues at the interface are shown as sticks. **B**) NBD1/ICL4 (left) and NBD2/ICL2 (right) in zCFTR. The coordinates for zCFTR correspond to the last frame of the ATP-bound 1 μs trajectory described in (10).

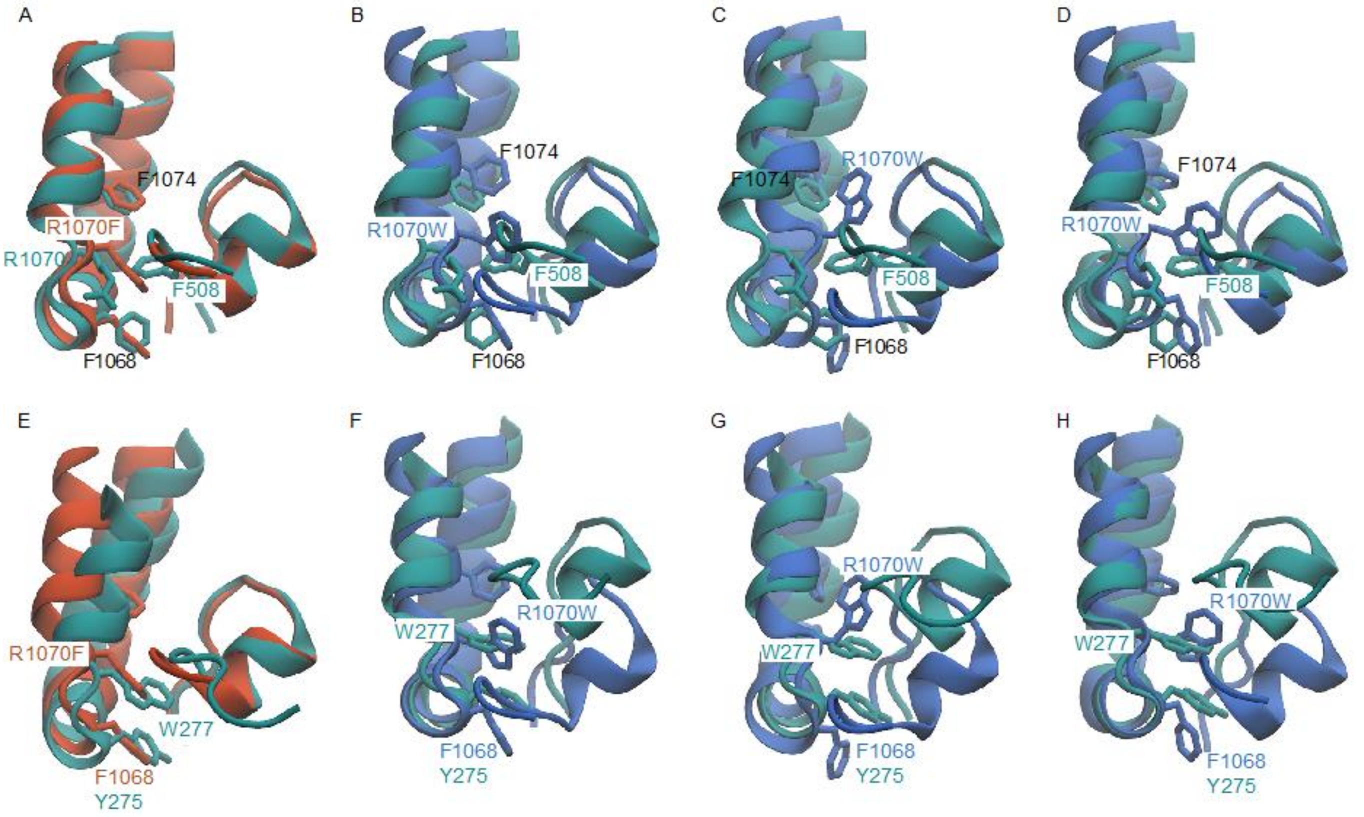


***Figure S9*. Comparison of the F508del/R1070F and F508del/R1070W cluster centers with the hCFTR ICL4/NBD1 interface**

**A-D**. Superimposition of the hCFTR ICL4/NBD1 interface (PDB ID 6MSM, (9). Teal cartoons) with the center of cluster 1 (**A**) from the F508del/R1070F system (orange cartoons) and the center of cluster 1 (**B**), cluster 2 (**C**), and cluster 3 (**D**) from the F508del/R1070W system (blue cartoons). Residues F1068, R1070X, F1074 and F508 are shown as sticks. **E-H**. Superimposition of the hCFTR ICL2/NBD2 interface (PDB ID 6MSM, (9). Teal cartoons) with the center of cluster 1 (**E**) from the F508del/R1070F system (orange cartoons) and the center of cluster 1 (**F**), cluster 2 (**G**), and cluster 3 (**H**) from the F508del/R1070W system (blue cartoons). Residues Y275 and W277 of hCFTR ICL2 and F1068, R1070X and F1074 from the simulation systems are shown as sticks.

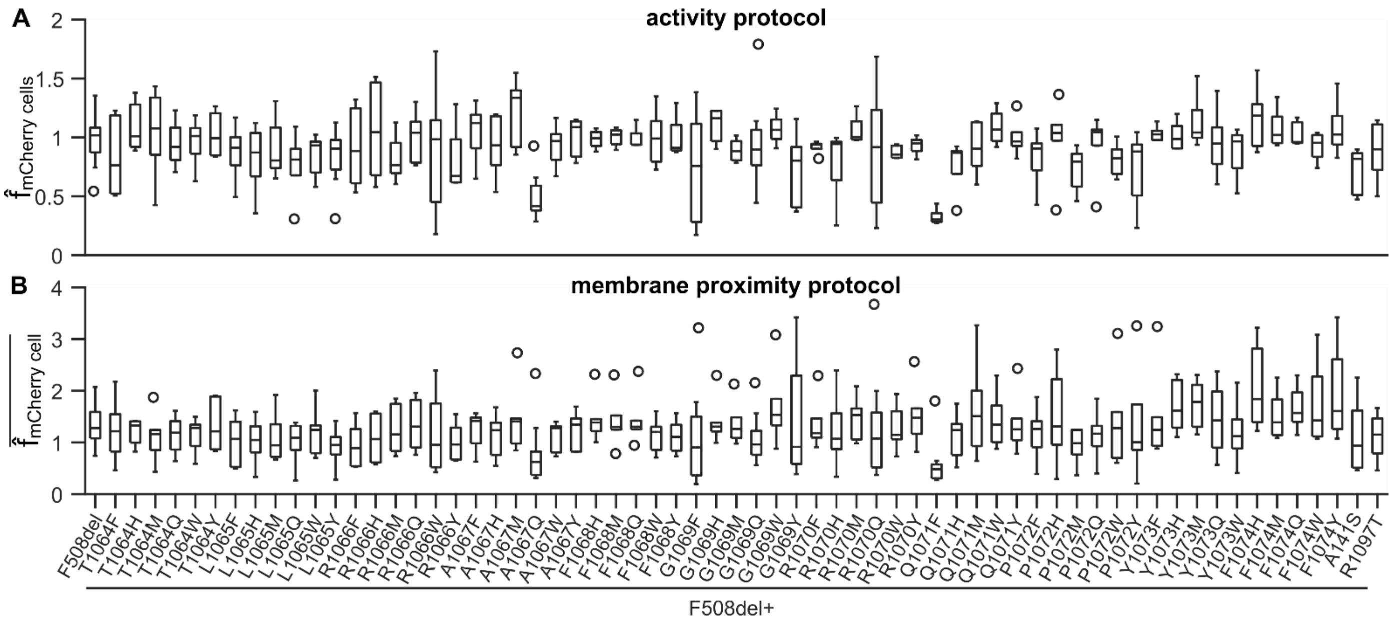


***Figure S10*** **Boxplots of the normalized mCherry fluorescence intensity**

The normalized mCherry fluorescence intensity for each variant was determined twice on each plate, as images were analysed using both the activity and the membrane proximity protocols. **A**. For measurements obtained in the CFTR activity protocol, the normalized mCherry fluorescence, $\hat{f}_{mCherry\ cells}$, was determined by normalizing the average mCherry fluorescence inside the cell selection for cells expressing each variant to that inside the cell selection of WT CFTR expressing cells on the same plate. **B**. In the membrane proximity protocol, the mCherry fluorescence, $f_{mCherry\ cell}$, was obtained for each cell separately, after which it was normalized to the median $f_{mCherry\ cell}$ of cells expressing WT CFTR on the same plate. This results in $\hat{f}_{mCherry\ cell}$ of which the plate means were taken for this boxplot (denoted as $\overline{\hat{f}_{mCherry\ cell}}$).

***Table S11*** **Descriptive statistics: normalized mCherry fluorescence intensity**

On every plate, the mean mCherry fluorescence intensity for each variant was determined for measurements obtained in the CFTR activity protocol and the membrane proximity protocol.

| | activity | | | | membrane proximity | | | |
|---|---|---|---|---|---|---|---|---|
| | **N** | **M** | **Mdn** | **SD** | **N** | **M** | **Mdn** | **SD** |
| WT | 20 | 1.00 | 1.00 | 0.00 | 23 | 1.33 | 1.25 | 0.19 |
| F508del | 19 | 0.98 | 1.02 | 0.18 | 22 | 1.35 | 1.28 | 0.38 |
| F508del + T1064F | 6 | 0.83 | 0.76 | 0.32 | 6 | 1.24 | 1.22 | 0.59 |
| F508del + T1064H | 5 | 1.09 | 1.01 | 0.21 | 6 | 1.21 | 1.33 | 0.25 |
| F508del + T1064M | 5 | 1.05 | 1.08 | 0.39 | 6 | 1.12 | 1.16 | 0.48 |
| F508del + T1064Q | 7 | 0.94 | 0.92 | 0.18 | 8 | 1.15 | 1.19 | 0.35 |
| F508del + T1064W | 5 | 0.96 | 1.01 | 0.21 | 6 | 1.15 | 1.28 | 0.34 |
| F508del + T1064Y | 5 | 1.03 | 0.99 | 0.19 | 6 | 1.32 | 1.22 | 0.48 |
| F508del + L1065F | 5 | 0.87 | 0.91 | 0.24 | 6 | 1.03 | 1.07 | 0.46 |
| F508del + L1065H | 5 | 0.83 | 0.87 | 0.29 | 6 | 1.02 | 1.05 | 0.45 |
| F508del + L1065M | 5 | 0.91 | 0.80 | 0.26 | 6 | 1.09 | 0.94 | 0.48 |
| F508del + L1065Q | 5 | 0.77 | 0.81 | 0.28 | 6 | 1.00 | 1.09 | 0.41 |
| F508del + L1065W | 5 | 0.84 | 0.93 | 0.18 | 6 | 1.22 | 1.24 | 0.46 |
| F508del + L1065Y | 10 | 0.83 | 0.90 | 0.23 | 11 | 0.94 | 0.95 | 0.30 |
| F508del + R1066F | 6 | 0.91 | 0.88 | 0.32 | 6 | 0.95 | 0.89 | 0.41 |
| F508del + R1066H | 5 | 1.06 | 1.04 | 0.42 | 6 | 1.08 | 1.07 | 0.45 |
| F508del + R1066M | 5 | 0.82 | 0.77 | 0.20 | 6 | 1.24 | 1.15 | 0.47 |
| F508del + R1066Q | 5 | 0.99 | 1.04 | 0.22 | 6 | 1.34 | 1.31 | 0.48 |
| F508del + R1066W | 9 | 0.87 | 0.98 | 0.48 | 11 | 1.13 | 0.95 | 0.70 |
| F508del + R1066Y | 5 | 0.81 | 0.67 | 0.28 | 6 | 1.01 | 0.96 | 0.37 |
| F508del + A1067F | 6 | 1.05 | 1.12 | 0.24 | 7 | 1.23 | 1.42 | 0.35 |
| F508del + A1067H | 6 | 0.92 | 0.93 | 0.25 | 7 | 1.10 | 1.24 | 0.41 |
| F508del + A1067M | 5 | 1.21 | 1.34 | 0.30 | 7 | 1.44 | 1.41 | 0.63 |
| F508del + A1067Q | 10 | 0.48 | 0.42 | 0.20 | 11 | 0.77 | 0.62 | 0.59 |
| F508del + A1067W | 5 | 0.93 | 0.97 | 0.18 | 6 | 1.13 | 1.28 | 0.29 |
| F508del + A1067Y | 5 | 1.00 | 1.09 | 0.17 | 6 | 1.25 | 1.34 | 0.35 |
| F508del + F1068H | 5 | 0.99 | 0.99 | 0.08 | 6 | 1.46 | 1.39 | 0.45 |
| F508del + F1068M | 5 | 1.00 | 1.02 | 0.08 | 6 | 1.41 | 1.30 | 0.50 |
| F508del + F1068Q | 5 | 0.99 | 0.94 | 0.09 | 6 | 1.43 | 1.30 | 0.49 |
| F508del + F1068W | 7 | 0.99 | 0.99 | 0.22 | 8 | 1.13 | 1.20 | 0.31 |
| F508del + F1068Y | 4 | 1.00 | 0.91 | 0.20 | 6 | 1.11 | 1.11 | 0.33 |
| F508del + G1069F | 10 | 0.73 | 0.76 | 0.45 | 10 | 1.11 | 0.91 | 0.90 |
| F508del + G1069H | 5 | 1.10 | 1.16 | 0.15 | 6 | 1.42 | 1.30 | 0.45 |
| F508del + G1069M | 5 | 0.89 | 0.88 | 0.10 | 6 | 1.37 | 1.26 | 0.42 |
| F508del + G1069Q | 10 | 0.95 | 0.90 | 0.36 | 11 | 1.07 | 0.96 | 0.46 |
| F508del + G1069W | 5 | 1.07 | 1.06 | 0.12 | 6 | 1.70 | 1.53 | 0.75 |
| F508del + G1069Y | 5 | 0.72 | 0.80 | 0.33 | 5 | 1.46 | 0.91 | 1.24 |
| F508del + R1070F | 5 | 0.92 | 0.94 | 0.06 | 6 | 1.35 | 1.18 | 0.49 |
| F508del + R1070H | 5 | 0.78 | 0.94 | 0.31 | 6 | 1.22 | 1.07 | 0.70 |
| F508del + R1070M | 5 | 1.06 | 1.00 | 0.12 | 6 | 1.47 | 1.53 | 0.41 |
| F508del + R1070Q | 11 | 0.87 | 0.92 | 0.48 | 11 | 1.29 | 1.08 | 0.96 |
| F508del + R1070W | 5 | 0.87 | 0.85 | 0.06 | 6 | 1.27 | 1.15 | 0.43 |
| F508del + R1070Y | 5 | 0.93 | 0.95 | 0.08 | 6 | 1.53 | 1.48 | 0.59 |
| F508del + Q1071F | 11 | 0.32 | 0.30 | 0.05 | 12 | 0.55 | 0.48 | 0.41 |
| F508del + Q1071H | 5 | 0.77 | 0.87 | 0.22 | 6 | 1.14 | 1.24 | 0.44 |
| F508del + Q1071M | 5 | 0.92 | 0.90 | 0.23 | 6 | 1.64 | 1.51 | 0.93 |
| F508del + Q1071W | 5 | 1.09 | 1.07 | 0.15 | 6 | 1.43 | 1.34 | 0.52 |
| F508del + Q1071Y | 5 | 1.00 | 0.97 | 0.16 | 6 | 1.37 | 1.25 | 0.57 |
| F508del + P1072F | 5 | 0.83 | 0.90 | 0.24 | 6 | 1.18 | 1.26 | 0.50 |
| F508del + P1072H | 6 | 0.98 | 1.04 | 0.33 | 7 | 1.52 | 1.31 | 0.86 |
| F508del + P1072M | 5 | 0.73 | 0.80 | 0.19 | 6 | 1.41 | 0.98 | 1.35 |
| F508del + P1072Q | 7 | 0.94 | 1.05 | 0.25 | 8 | 1.13 | 1.17 | 0.42 |
| F508del + P1072W | 5 | 0.81 | 0.82 | 0.14 | 6 | 1.43 | 1.28 | 0.91 |
| F508del + P1072Y | 5 | 0.73 | 0.88 | 0.32 | 6 | 1.34 | 1.00 | 1.06 |
| F508del + Y1073F | 5 | 1.03 | 1.03 | 0.07 | 6 | 1.51 | 1.24 | 0.89 |
| F508del + Y1073H | 5 | 1.01 | 0.99 | 0.12 | 6 | 1.69 | 1.61 | 0.53 |
| F508del + Y1073M | 5 | 1.13 | 1.04 | 0.23 | 6 | 1.74 | 1.79 | 0.48 |
| F508del + Y1073Q | 7 | 0.96 | 0.95 | 0.26 | 8 | 1.44 | 1.43 | 0.65 |
| F508del + Y1073W | 5 | 0.87 | 0.97 | 0.22 | 6 | 1.19 | 1.12 | 0.59 |
| F508del + F1074H | 5 | 1.15 | 1.18 | 0.27 | 6 | 2.05 | 1.84 | 0.80 |
| F508del + F1074M | 5 | 1.08 | 1.02 | 0.17 | 6 | 1.51 | 1.39 | 0.45 |
| F508del + F1074Q | 5 | 1.03 | 0.96 | 0.10 | 6 | 1.66 | 1.57 | 0.42 |
| F508del + F1074W | 5 | 0.92 | 0.95 | 0.12 | 6 | 1.73 | 1.43 | 0.79 |
| F508del + F1074Y | 5 | 1.08 | 1.03 | 0.23 | 6 | 1.93 | 1.61 | 0.91 |
| F508del + A141S | 5 | 0.71 | 0.82 | 0.20 | 6 | 1.12 | 0.94 | 0.70 |

***Text S12Mathematical model***

HEK-293 cells were modelled as 8.9 µm-radius spheres in the presence of intra- and extracellular $Cl^-$, $K^+$, and $I^-$ at 28 °C. To account for the effect of filopodia on the membrane surface area of HEK-293 cells (1), the membrane surface area ($A_m$) of the modelled HEK-293 cells was adjusted by adding 50% to the value calculated from their radius ($r$); $A_m = 4\pi r^2 + \frac{1}{2}(4\pi r^2)$. The volume of the cells was modelled as $V_{cell} = \frac{4}{3}\pi \cdot r^3$. Changes in the system were modelled at time intervals of 0.2 ms. The free parameters in the model were $G_{CFTR-Cl}$, and $V_m$. The maximal $G_{CFTR-Cl}$ (G in nS) and the membrane potential before addition of iodide ($V_m$ in mV) were used as assay readouts to quantify CFTR function***Table*** . In the model, R, T and F have their usual meaning. R is the ideal gas constant (8.314 $J\cdot K^{-1}\cdot mol^{-1}$), T is the absolute temperature in Kelvin, and F is Faraday's constant (96485.332 $C\cdot mol^{-1}$).

Initial concentrations of $Cl^-$, $K^+$ and $I^-$

In the model, time point 0 represents the moment of iodide addition to the extracellular medium. At this moment CFTR activity has reached a steady-state, and $[Cl^-]_{in}$ is assumed to have equilibrated with $[Cl^-]_{out}$. The initial $Cl^-$, $K^+$ and $I^-$ concentrations at time point 0 are:

| | |
|---|---|
| $[Cl^-]_{out}$ | 117.1 mM (corresponding to the extracellular $Cl^-$ concentration after $I^-$ addition) |
| $[Cl^-]_{in}$ | $[Cl^-]_{out}/e^{\left(\frac{V_m}{RT/z_iF}\right)}$ where $[Cl^-]_{out}$ is 152 mM corresponding to the extracellular $Cl^-$ concentration before $I^-$ addition ($z_i$ is the valency of ion $i$) |
| $[K^+]_{out}$ | 4.7 mM |
| $[K^+]_{in}$ | 100.0 mM |
| $[I^-]_{out}$ | 100.0 mM |
| $[I^-]_{in}$ | 0.0 mM |

Ionic currents and conductance

The Goldman-Hodgkin-Katz flux equation for an ion $i$ describes the ionic current ($I_i$ in $A\cdot m^{-2}$) across a cell membrane as a function of the membrane potential ($V_m$), the permeability of the membrane to ion $i$ ($p_i$), the valency of ion $i$ ($z_i$),and the concentrations of the ion inside ($[i]_{in}$) and outside ($[i]_{out}$) of the cell:

$$I_i = \frac{p_i {z_i}^2 F^2}{RT} V_m \left( \frac{[i]_{in} - [i]_{out} e^{-z_i F V_m/RT}}{1 - e^{-z_i F V_m/RT}} \right)$$

In symmetrical solutions, where both extracellular and intracellular concentrations of ion $i$ are $[i]_{sym}$ (see Appendix A in (2)) the conductance ($G_i$) for ion $i$ can be expressed as $G_i = \frac{p_i {z_i}^2 F^2 [i]_{sym}}{RT}$, which can be rearranged as $\frac{G_i}{[i]_{sym}} = \frac{p_i {z_i}^2 F^2}{RT}$, making it possible to express whole-cell ionic currents of ion $i$ ($I_i$) as follows:

$$I_i = \frac{G_i}{[i]_{sym}} V_m \left( \frac{[i]_{in} - [i]_{out} e^{-z_i F V_m/RT}}{1 - e^{-z_i F V_m/RT}} \right)$$

Using the maximal conductance values in symmetrical solutions of 140 mM, whole-cell currents for $Cl^-$, $K^+$ and $I^-$ were predicted for our experimental conditions. $K^+$ currents ($I_K$) in HEK-293 cells are mediated by endogenous potassium channels. An endogenous leak conductance for $K^+$ ($G_{leak-K}$) was set to 2.5 nS (3) to predict $K^+$-mediated currents ($I_K$). The CFTR-mediated $Cl^-$ conductance ($G_{CFTR-Cl}$) was estimated by fitting the experimental data to the model. The parameter was constrained between 0 and 300 nS to avoid unphysiological values.

The permeability and conductance of WT CFTR for $Cl^-$ are higher than those for $I^-$; the permeability to $I^-$ over the permeability to $Cl^-$ ($p_I/p_{Cl}$) is 0.83 (4). Because $Cl^-$ and $I^-$ ions have the same valency ($z_{Cl} = z_I$), in symmetrical solutions we can expect the relationship between CFTR-mediated $Cl^-$ and $I^-$ conductance ($G_{CFTR-I}/G_{CFTR-Cl}$) to be similar. Although this is obviously a simplification, the CFTR-mediated $I^-$ conductance ($G_{CFTR-I}$) was set to be proportional to the CFTR-mediated $Cl^-$ conductance ($G_{CFTR-I} = 0.83 \cdot G_{CFTR-Cl}$). A non-CFTR related transient anion conductance ($G_{trans}$) observed upon addition of $I^-$, had to be added to the model to describe the experimental data accurately (5). We hypothesized that endogenous anion permeabilities of the HEK-293 cells underlie the transient conductance, triggered upon I⁻ addition. The time course of $G_{trans}$ is described in the model as a single exponential decay characterized by a time constant ($\tau_{trans}$). For this transient conductance too, its whole-cell current was predicted at each time point and it was added to the estimated CFTR-mediated Cl⁻ and I⁻ currents ($I_{Cl}$ and $I_I$).

$$I_K = \frac{G_{leak-K}}{[K^+]_{sym}} V_m \left( \frac{[K^+]_{in} - [K^+]_{out} e^{-z_i F V_m / RT}}{1 - e^{-z_i F V_m / RT}} \right)$$

$$I_{Cl} = \frac{G_{CFTR-Cl}}{[Cl^-]_{sym}} V_m \left( \frac{[Cl^-]_{in} - [Cl^-]_{out} e^{-z_i F V_m / RT}}{1 - e^{-z_i F V_m / RT}} \right) + \frac{G_{trans} e^{-t/\tau_{trans}}}{[Cl^-]_{sym}} V_m \left( \frac{[Cl^-]_{in} - [Cl^-]_{out} e^{-z_i F V_m / RT}}{1 - e^{-z_i F V_m / RT}} \right)$$

$$I_I = \frac{0.83 \cdot G_{CFTR-Cl}}{[I^-]_{sym}} V_m \left( \frac{[I^-]_{in} - [I^-]_{out} e^{-z_i F V_m / RT}}{1 - e^{-z_i F V_m / RT}} \right) + \frac{0.83 \cdot G_{trans} e^{-t/\tau_{trans}}}{[I^-]_{sym}} V_m \left( \frac{[I^-]_{in} - [I^-]_{out} e^{-z_i F V_m / RT}}{1 - e^{-z_i F V_m / RT}} \right)$$

Time dependence of intracellular concentrations of Cl⁻, K⁺ and I⁻

The molar ion flux per second for ion $i$ is given by $\frac{I_i}{z_i F}$. The intracellular concentrations at subsequent timepoints ($[i]_{in}(t+1)$) were approximated as follows, using the intracellular concentration of ion $i$ at timepoint t ($[i]_{in}(t)$) and the estimated ionic current of ion $i$:

$$[i]_{in}(t+1) = [i]_i(t) + \mathrm{d}t \frac{-\left(\frac{I_i}{z_i F}\right)}{V_{cell}}$$

Membrane potential

The membrane capacitance determines the rate at which the membrane potential changes in response to the charge that moves across the membrane. A membrane capacitance of 1 $\mu F \cdot cm^{-2}$, typically used for biological membranes (6), was used to estimate the membrane capacitance of the cell. The membrane capacitance ($C_m$) is constant over time, and in our model, the net ionic current is determined by a K⁺ leak current, and Cl⁻ and I⁻ currents with CFTR and non-CFTR-mediated components ($I_{ion} = I_K + I_{Cl} + I_I$). The new membrane potential ($V_{m(t+1)}$) at subsequent time points was estimated as follows:

$$V_{m(t+1)} = V_m(t) + dt \frac{-(I_K + I_{Cl} + I_I)}{C_m}$$

The membrane potential estimates were constrained between –30 mV and –90 mV to avoid unphysiological values.

The proportion of anion-bound and anion-free YFP(H148Q/I152L)

The anion-binding site on the YFP(H148Q/I152L) chromophore, can be unoccupied, bound to I⁻ or bound to Cl⁻. The relative proportions of binding sites occupied by I⁻ and Cl⁻ depend on their intracellular concentrations and the affinities of the halides for the binding site. The binding affinities of I⁻ and Cl⁻ to YFP(H148Q/I152L) are 1.9 mM and 85 mM, respectively (7). The intracellular ion concentrations ($[I^-]$ and $[Cl^-]$) at every simulated time point are determined as described in the section above. To estimate the proportion of I⁻ bound ($P_I$), Cl⁻ bound ($P_{Cl}$), and anion-free YFP(H148Q/I152L) ($P_{free}$) we used the following equations:

$$P_I = \frac{[I^-]}{K_I\left(1 + \frac{K_{Cl}}{[Cl^-]}\right) + [I^-]}$$

$$P_{Cl} = \frac{[Cl^-]}{K_{Cl}\left(1 + \frac{[I^-]}{K_I}\right) + [Cl^-]}$$

$$P_{free} = 1 - (P_I + P_{Cl})$$

Because only anion-free YFP(H148Q/I152L) is fluorescent in our experimental conditions (8), we can fit the $P_{free}$ predictions from the model to observed experimental fluorescence measurements to estimate the free parameters. To better relate to the fluorescence quenching time course to $P_{free}$, $P_{free}$ was normalized to $P_{free}$ at time point zero $(t0)$.

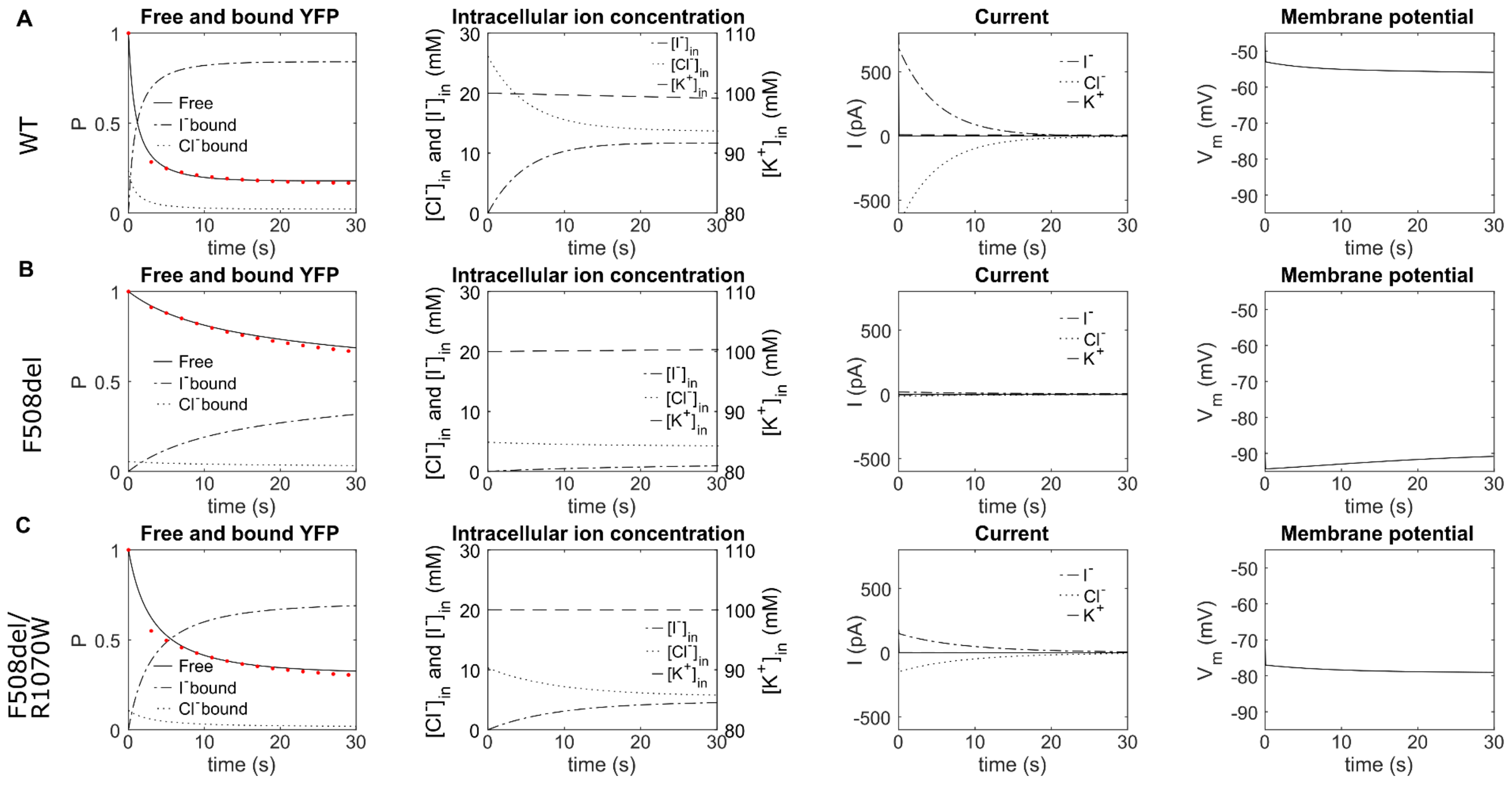


***Figure S13*** **Fitting of the fluorescence quenching to a simple mathematical model**

HEK-293 cells were transfected with pIRES2-mCherry-YFPCFTR containing either WT CFTR (**A**), F508del-CFTR (**B**) or F508del/R1070W-CFTR (**C**). CFTR was activated with 10 µM forskolin and 230 s allowed for activation to reach steady-state before $I^-$ addition at time point 0. The first column of panels displays the measured normalized YFP(H148Q/I152L) fluorescence (filled red circles). These measurements were used to obtain fit parameters to model the proportions of anion-free, $Cl^-$ bound and $I^-$ bound YFP(H148Q/I152L). To relate to the normalized fluorescence quenching time course (filled red circles), the proportion of free YFP(H148Q/I152L) was also normalized to the time point just before addition of $I^-$. Other panels show the modelled intracellular ion concentrations and transmembrane ion currents carried by $Cl^-$, $I^-$ and $K^+$and the modelled membrane potential.
